# SCITO-seq: single-cell combinatorial indexed cytometry sequencing

**DOI:** 10.1101/2020.03.27.012633

**Authors:** Byungjin Hwang, David S. Lee, Whitney Tamaki, Yang Sun, Anton Ogorodnikov, George Hartoularos, Aidan Winters, Yun S. Song, Eric D. Chow, Matthew H. Spitzer, Chun Jimmie Ye

## Abstract

The development of DNA-barcoded antibodies to tag cell-surface molecules has enabled the use of droplet-based single cell sequencing (dsc-seq) to profile the surface proteomes of cells. Compared to flow and mass cytometry, the major limitation of current dsc-seq-based workflows is the high cost associated with profiling each cell, thus precluding its use in applications where millions of cells are required. Here, we introduce SCITO-seq, a new workflow that combines combinatorial indexing and commercially available dsc-seq to enable cost-effective cell surface proteomic sequencing of greater than 10^5^ cells per microfluidic reaction. We demonstrate SCITO-seq’s feasibility and scalability by profiling mixed species cell lines and mixed human T and B lymphocytes. To further demonstrate its applicability, we show comparable cellular composition estimates in peripheral blood mononuclear cells obtained with SCITO-seq and mass cytometry. SCITO-seq can be extended to include simultaneous profiling of additional modalities such as transcripts and accessible chromatin or tracking of experimental perturbations such as genome edits or extracellular stimuli.

## Main

The use of DNA to barcode physical compartments and tag intracellular and cell-surface molecules has enabled the use of sequencing to efficiently profile the molecular properties of thousands of cells simultaneously. While initially applied to measuring the abundances of RNA^1,2^ and identifying regions of accessible DNA^3^, recent developments in DNA-tagged antibodies have created new opportunities to use sequencing to measure the abundances of cell surface^4,5^ and intracellular proteins^6^.

Sequencing DNA-tagged antibodies is particularly useful for profiling cells whose identity and function have long been determined by the expression of cell surface proteins (e.g. immune cells) and has several advantages over flow and mass cytometry. First, the number of cell surface proteins that can be measured by DNA-tagged antibodies is exponential to the number of bases in the tag. In theory, all cell surface proteins with available antibodies can be targeted and in practice, panels targeting hundreds of proteins are now commercially available^4,7^. This contrasts with cytometry where the number of proteins targeted is limited by the overlap in the emission spectrums of fluorophores (flow: 4-48) or the number of unique masses of metal isotopes that can be chelated by commercial polymers (CYTOF: ~50)^8,9^. Second, sequencing-based proteomics can readily read out all antibody-derived tags (ADTs) with one reaction instead of subsequent rounds of signal separation and detection, significantly reducing the time and sample input for profiling large panels and obviating the need for fixation. Third, additional molecules can be profiled within the same cell enabling multimodal profiling of cell surface proteins along with the immune repertoire, transcriptome^4^, and potentially the epigenome. Finally, sequencing is amenable to encoding orthogonal experimental information using additional DNA barcodes (either inline or distributed) creating opportunities for large-scale multiplexed screens that barcode cells using natural variation^10^, synthetic sequences^11,12^, or sgRNAs^13,14^.

Compared to flow and mass cytometry, the major limitation in sequencing-based single-cell proteomics^4,7^ is the high cost associated with profiling each cell, thus precluding its use across population cohorts or large-scale screens where millions of cells would need to be profiled. Like other single-cell sequencing assays, total cost per cell for proteomic sequencing is divided between cost associated with library construction and the cost for sequencing the library. Because the number of protein molecules per cell is 2-6 orders of magnitude higher than RNA^15^ and the use of targeting antibodies limits the number of features measured per cell, sequencing ADTs will result in more unique molecules than RNA given the same depth of sequencing per cell. However, the costs associated with commercially available microfluidics-based single cell library construction^16^ and conjugation of modified DNA sequences to antibodies^4^ are high. Thus, for single-cell proteomic sequencing to be a compelling strategy for high dimensional phenotyping of millions of cells, there is a major need to develop a workflow that minimizes library and antibody preparation costs.

Here, we introduce <u>s</u>ingle cell <u>c</u>ombinatorial <u>i</u>ndexed cy<u>to</u>metry by <u>seq</u>uencing (SCITO-seq), a single cell proteomics workflow that combines split-pool indexing and droplet-based sequencing. Our approach is based on the key insight that the large number of droplets produced by microfluidic workflows (e.g. ~10^5^ for 10X Genomics^16^) can be used as a second round of physical compartments for single-cell combinatorial indexing (SCI)^17–20^ resulting in a simple and cost-effective two-step procedure for library construction. While two-step SCI workflows have been recently described for ATAC-seq^21^ and RNA-seq^22^, direct implementation of combinatorial indexing for protein sequencing has not been reported. For SCITO-seq, we introduce a strategy using universal conjugation followed by pooled hybridization to minimize the cost for generating large panels of DNA-tagged antibodies. Tagged antibodies are then used to stain cells in individual pools prior to high-concentration loading utilizing commercially available microfluidics. After ADT library construction and sequencing, protein expression profiles for cells simultaneously encapsulated in a single drop are resolved by the combinatorial index of pool and droplet barcodes. Compared to other single cell sequencing workflows, loading cells at high concentrations and sequencing ADTs associated with a limited number of cell surface proteins reduce the library construction and sequencing costs per cell, respectively. We demonstrate the feasibility and scalability of SCITO-seq in mixed species and mixed individual experiments profiling 10^4^-10^5^ cells per microfluidic reaction. We further illustrate an application of SCITO-seq by profiling up to 10^5^ peripheral blood mononuclear cells using a panel of 28 antibodies in one microfluidic reaction, producing results comparable to mass cytometry (CyTOF) while achieving >10-fold increase in throughput compared to standard sequencing workflows. Finally, we demonstrate that targeted sequencing of ADTs using SCITO-seq can recover the same cell clusters at very low sequencing depths per cell. SCITO-seq can be integrated with existing workflows for multimodal profiling of transcripts^22^ and accessible chromatin^21^ and can be a compelling platform for obtaining rich phenotyping data from high-throughput screens of genetic and extracellular perturbations.

As microfluidic cell loading is governed by a Poisson distribution, the major limitation of standard droplet-based single cell sequencing (dsc-seq) workflows is ensuring encapsulation of single cells to reduce the number of collisions. This results in suboptimal cell recovery, reagent usage, and inflated library construction costs. For the 10X Genomics microfluidic platform, Poisson loading at concentrations of 2×10^3^-2×10^4^ cells results in a cell recovery rate (CRR) of 50-60%^16,22^ and collision rates of 1-10% (**Supplementary Fig. 1**). However, greater than 97%-82% of droplets do not contain a cell, leading to wasted reagents (**Supplementary Fig. 2 and 3**). One approach to decrease the library preparation cost and increase the sample and cell throughput of dsc-seq is to “barcode” samples using either natural genetic variants^10,23,24^ or synthetic DNA molecules^11,12,25^ prior to pooled loading at 5×10^4^-8×10^4^ cells, reducing the proportion of droplets without a cell to ~65%-45%. Because simultaneous encapsulation of cells within a droplet can be detected by the co-occurrence of different sample barcodes (e.g. genetic variant or synthetic DNA tags) with the same droplet barcode (DBC), sample multiplexing increases the number of singlets recovered per microfluidic reaction while maintaining a low effective collision rate that is tunable by the number of sample barcodes. However, as collision events can only be detected but not resolved into usable single-cell data, the maximum number of cells loaded to minimize cost is ultimately limited by the overhead cost incurred for sequencing collided droplets.

Single-cell combinatorial indexing (SCI) is an alternative, scalable approach to control the collision rate of single-cell sequencing by labeling subsequent rounds of physical compartmentalization with DNA barcodes. While standard SCI approaches require more than two rounds of combinatorial indexing to sequence 10^5^-10^6^ cells^17–20^, recent advances utilizing droplet-based microfluidics for combinatorial indexing have enabled simplified two-round workflows to achieve the same throughput^21,22^. For applications where only a set of targeted markers are needed, such as high-throughput screens and clinical biomarker profiling, current SCI workflows profiling the entire epigenome or transcriptome per cell would likely result in prohibitively high sequencing costs.

Here we propose a simple experimental workflow, SCITO-seq, which combinatorically indexes single cells using DNA-tagged antibodies^4^ and microfluidic droplets to enable cost-effective profiling of cell-surface proteins scalable to 10^5^-10^6^ cells (**Fig. 1a**). First, each antibody is conjugated with specific amine modified oligonucleotide sequences (antibody handle, 20bp) which enables pooled hybridization to minimize the costs associated with generating large numbers of DNA-tagged antibodies. Second, equal amounts of antibodies are combined and aliquoted before the addition of an oligo pool containing both a compound barcode to delineate the antibody and the pool (Ab+PBC), as well as a complementary sequence to the antibody handle for hybridization to its respective antibody (**Fig. 1b**). Third, cells are allocated into pools and stained with pool-specific antibodies. Fourth, the stained cells are combined and loaded at concentrations tunable to the targeted collision rate followed by processing using a commercially available dsc-seq platform to generate a sequencing library. Finally, after sequencing only the antibody derived tags (ADTs), the surface protein expression profiles of multiple encapsulated cells (multiplets) within a droplet can be resolved into usable single-cell protein expression profiles by the combinatorial index of Ab+PBC and DBC.

**Figure 1:**
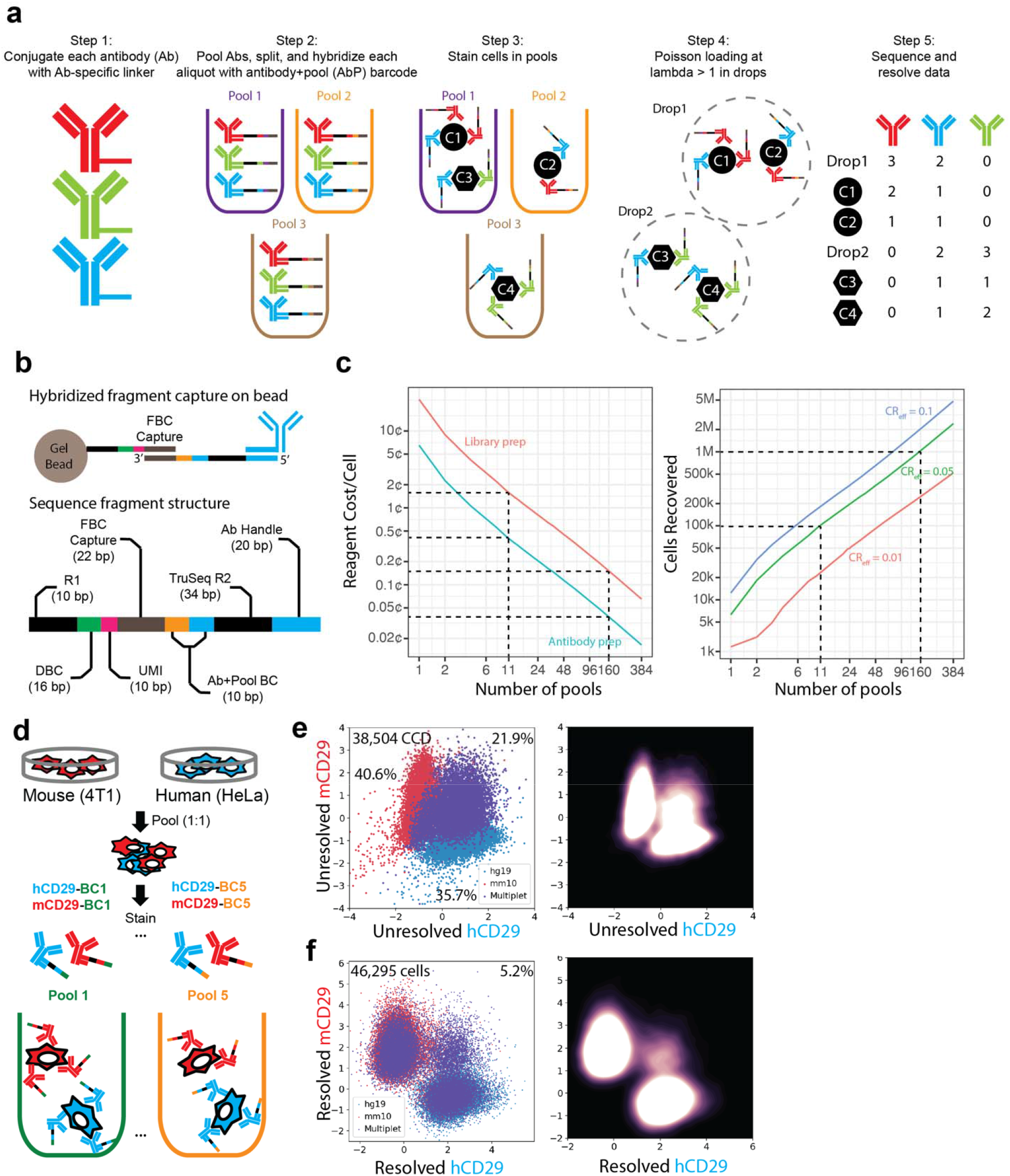
Design of SCITO-seq and mixed-species proof-of-concept experiment. **(a)** SCITO-seq workflow. Antibodies are first each conjugated with a unique antibody barcode (red, blue and green) and hybridized with an oligo containing the compound antibody and pool barcodes (Ab+Pool BC: [red, blue, green] × [purple, orange, brown]). Cells are split and stained with pool-specific antibodies per pool. Stained cells are combined and loaded for droplet-based sequencing at high concentrations. Cells are resolved from the resulting data using the combinatorial index of Ab+Pool BC and droplet barcodes. **(b)** A detailed structure of the SCITO-seq fragment produced. The primary universal oligo is an antibody specific hybridization handle. The secondary oligo consists of the reverse complement sequence to the handle followed by a TruSeq adaptor (black), the compound Ab+Pool barcode (orange+blue), and the 10x 3’v3 feature barcode sequence (FBC, gray). The Ab+Pool barcode and the droplet barcode (DBC) form a combinatorial index unique to each cell. **(c)** Cost savings and collision rate analysis. As the number of pools increases, total library and DNA-barcoded antibody construction costs drop (left) while the number of cells recovered increases (right). Number of cells recovered as a function of the number of pools at three commonly accepted collision rates (1%, 5% and 10%) (right). **(d)** Mixed species (HeLa and 4T1) proof-of-concept experiment. HeLa and 4T1 cells are mixed and stained in five separate pools at a ratio of 1:1 with SCITO-seq antibodies barcoded with pool-specific barcodes. Scatter (left) and density (right) plots of **(e)** 38,504 unresolved cell-containing droplets (CCD) and **(f)** 46,295 resolved cells at a loading concentration of 10^5^ cells. Merged ADT counts are generated by summing all counts for each antibody across pools simulating standard workflows. Resolved data is obtained after assigning cells based on the combination of Ab+Pool and DBC barcodes.

Key to SCITO-seq is the recognition that because of Poisson loading, the fraction of droplets with increasing number of cells decreases exponentially. Thus, indexing cells using a small number of antibody pools will ensure that the combinatorial index (Ab+PBC and DBC) will identify a cell at low collision rates even at high loading concentrations. Theoretically, given *P* pools, *C* cells loaded, *D* droplets formed, the collision rate is given as 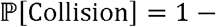 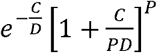 while the rate of empty droplets is given by 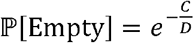 (**see Methods** for derivation). Our derivation of the collision rate differs from previously reported estimates derived from the classical birthday problem^22^, which did not account for higher order encapsulation of more than two cells within the same droplet. These closed form derivations of the collision and empty droplet rates are nearly identical to those obtained based on simulations (**Supplementary Fig. 4**). For example, in a commercially available microfluidic platform where 10^5^ droplets are formed, a loading concentration of 1.82×10^5^ cells (target recovery of 10^5^ cells) results in 84% of droplets containing at least one cell but only 4.4% of droplets containing greater than four cells (**Supplementary Fig. 2 and 3**). To yield 10^5^ resolved cells at a collision rate of 5% for this loading concentration, 11 antibody pools would be needed, resulting in an antibody preparation and library construction cost of 2¢/cell and savings of 17-fold over the standard workflow (**Fig. 1c**). At 160 pools and 5% collision rate, 10^6^ cells can be profiled in one microfluidic reaction with an average of 18.9 cells captured per droplet resulting in a total preparation cost of 0.2¢/cell and savings of 174-fold over the standard workflow (**Fig. 1c**).

To demonstrate the feasibility of SCITO-seq, we first performed a mixed species experiment by mixing human (HeLa) and mouse (4T1) cells, splitting into five aliquots, and staining each pool with anti-human CD29 (hCD29) and anti-mouse CD29 (mCD29) antibodies labeled with pool-specific barcodes (**Fig. 1d**). After washing unbound antibodies and mixing the five stained pools at equal proportions, 10^5^ cells were loaded for ADT library construction using the 10X Genomics 3’ V3 chemistry and the resulting library sequenced to recover 38,504 post-filtered cell-containing droplets (CCDs) at a depth of 2,909 reads/CCD. For comparison purposes, we also prepared and sequenced a library derived from the transcriptome at a depth of 25,844 reads/CCD. Merging ADTs for each antibody across pools to mimic standard single-cell proteomic profiling^4^, we detected 40.6% and 35.7% of CCDs with only mouse or human CD29 ADTs and 21.9% with CD29 ADTs from both species which we labeled as cross-species multiplets (**Fig. 1e**). Estimates from the analyses of transcriptomic data yielded similar results: 38.1% CCDs with mouse transcripts, 32.7% with human transcripts and 29.1% cross-species (**Supplementary Fig. 5**). Based on the frequencies of mouse and human singlets, an additional 22.0% of CCDs are estimated to be within-species multiplets and cannot be detected or resolved based on the merged ADTs for a total of 44% multiplets. Utilizing the combinatorial DBC and Ab+PBCs, the cross-species collision rate was reduced to 5.2% (**Fig. 1f**) and many within-species multiplets were also resolved (**Supplementary Fig. 6**) without significant pool to pool variation (**Supplementary Fig. 7)**. The ability to resolve cross and within-species multiplets results in a total of 46,295 cells profiled at an estimated collision rate of 11.0%, a 3.7-fold increase over standard workflows (12,500 droplets at 11% collision rate) (**Fig. 1f**). Similar results were observed for a two-pool design and lower loading concentration of 2×10^4^ cells, using universal conjugation achieving a 2.2-fold increase in throughput at a collision rate of 4.6% (**Supplementary Fig. 7, 8**, **9 and 10**). Our results are also consistent with an alternative two-pool design where four different Ab+PBC barcodes were directly conjugated onto the antibodies suggesting that both within and across pool contamination rates of secondary hybridization oligos are low (**Supplementary Fig. 11**).

We next sought to further assess the scalability of SCITO-seq and its applicability to resolve quantitative differences in cellular composition based on cell surface protein expression. We isolated primary CD4^+^ T cells and B cells from two donors and mixed them at a ratio of 5:1 for one donor and 1:3 for the second donor. To account for potential batch effects from SCITO-seq, we equally mixed donors prior to splitting and staining each of five pools with pool-barcoded CD4 and CD20 antibodies (**Fig. 2a**). Stained pools were mixed at equal ratios, loaded at 10^5^ and 2×10^5^ cells per well on a 10X Chromium system, processed with 3’V3 chemistry, and the resulting ADT and RNA libraries sequenced to recover 41,889 and 58,769 post-processing CCDs, respectively.

**Figure 2:**
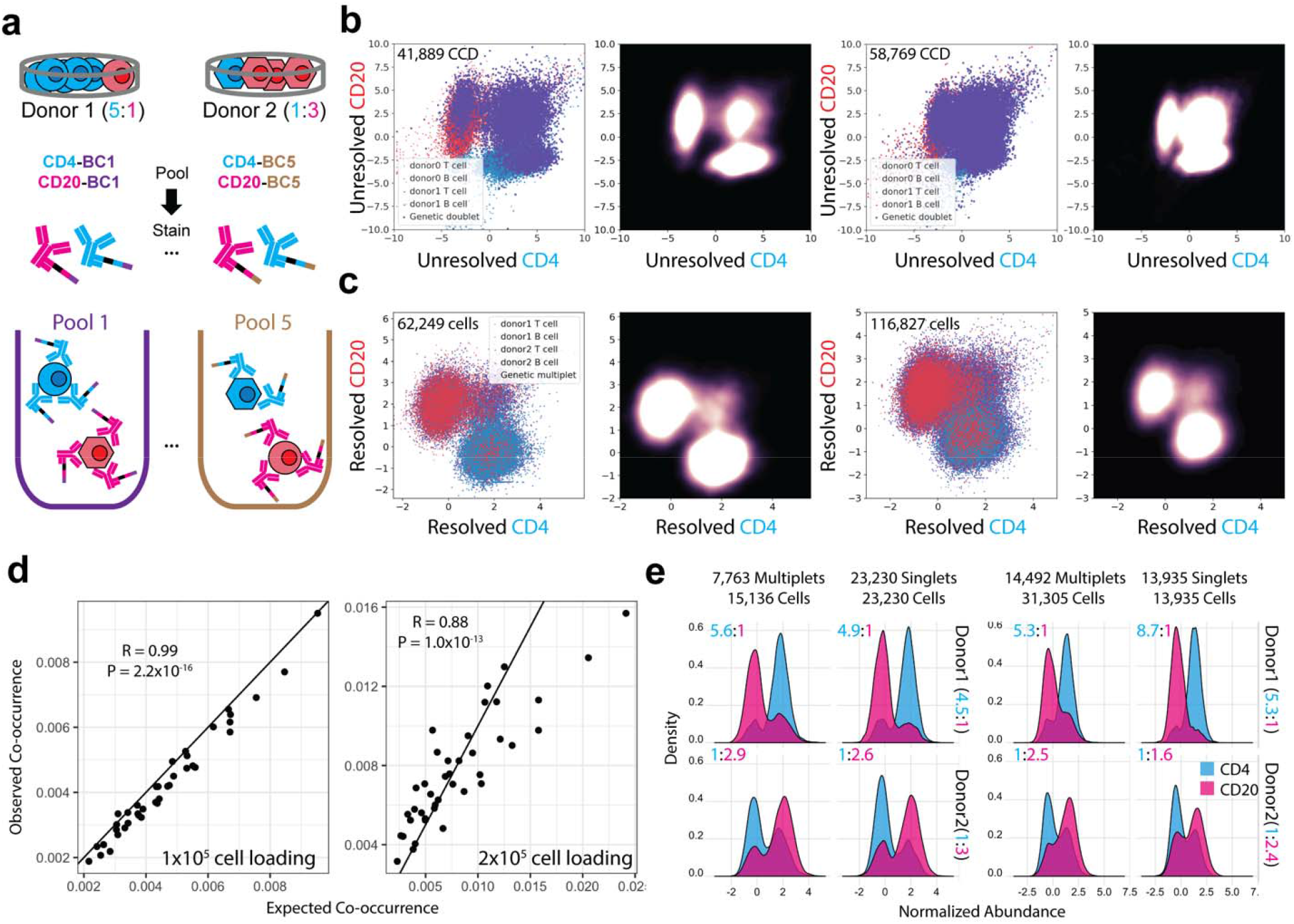
Demonstration of SCITO-seq in human donor experiment with significant increase in throughput of profiling proteins. **(a)** Schematic of human mixing experiment where different ratios of T and B cells (5:1 and 1:3) were mixed prior to splitting and indexing with five pools of CD4 and CD20 antibodies. Cell types are indicated by color (T: blue, B: red) while shapes indicate donors. Side by side scatter plot and density plots of **(b)** unresolved and **(c)** resolved cells for loading concentrations of 10^5^ (first two from the left) and 2×10^5^ (third and fourth from the left) cells. **(d)** Expected (x-axis) versus observed (y-axis) frequencies of co-occurrences between antibody pool barcodes for loading concentrations of 10^5^ (left) and 2×10^5^ (right) cells. Expected frequencies were calculated based on the frequencies of barcodes in pool singlets. **(e)** Distribution of the normalized UMI counts for each antibody in cells resolved from pool singlets and pool multiplets per donor. Distribution of the antibodies in pool multiplets shows expected prior mixture proportions and overlaps with the corresponding distribution in pool singlets.

Merging the ADT data across the five pools, CD4 and CD20 antibodies stained the expected cell types defined by the transcriptome and identified 25.7% (10^5^ cell loading) and 40% (2×10^5^ cell loading) cross cell-type multiplets consistent with the transcriptomic data (10^5^: 23.5%; 2×10^5^: 49.6%, **Fig. 2b**, **Supplementary Fig. 12**). We further used demuxlet^10^ to analyze genetic variants captured in the transcriptomic data to estimate 17.7% (10^5^ cell loading) and 30% (2×10^5^ cell loading) within cell-type multiplets that cannot be resolved by merged ADTs for a total multiplet rate of 43.4% and 70%. After resolving both cross and within cell-type multiplets using the combinatorial index of Ab+Pool and DBC with minimal batch effects (**Supplementary Fig. 13)**, we obtained 60,249 and 116,827 resolved cells with the frequency of cross cell-type collision events reduced from 25.7% to 8.9% (10^5^ cell loading) and from 40% to 14.3% (2×10^5^ cell loading), while the frequencies of single positives increased (**Fig. 2b** **and** **c**). Further, the observed co-occurrences of SCITO-seq antibodies from different pools were highly correlated with the expected co-occurrences (10^5^ cell loading: R = 0.99, P < 2.2×10^−16^; 2×10^5^ cell loading: R = 0.88, P < 1.0×10^−13^), suggesting that the encapsulation of multiple cells within droplets is not biased for specific pools or cell types (**Fig. 2d)**. These results demonstrate the scalability of SCITO-seq to profile 60,249 and 116,827 cells in one microfluidic channel at collision rates of 15.0% and 25.0%, effectively increasing the throughput by 3.3-fold and 4.0-fold over standard workflows at the same collision rates.

We next assessed if SCITO-seq can capture the unequal distribution of B and T cells from the two donors, especially from droplets that encapsulated multiple cells. For this analysis, we focused only on 39,955 (10^5^ cell loading) and 45,240 (2×10^5^ cell loading) droplets predicted to contain cells from one genetic background. Within droplets expressing only one antibody pool barcode (pool singlets), analysis of the frequencies of CD4^+^ and CD20^+^ cells (T:B_100K_: 4.9:1 and 1:2.6; T:B_200K_: 8.7:1 and 1:1.6) mirrored the expected proportion of T to B cells for each of the two donors and was consistent with estimates obtained from the transcriptomic data (**Fig. 2e**). Encouragingly, approximately the same frequency estimates were obtained in droplets expressing multiple pool (pool doublets) barcodes (T:B_100K_: 5.6:1 and 1:2.9; T:B_200K_: 5.3:1 and 1:2.5, **Fig. 2e**). These results highlight the ability of SCITO-seq not only to detect but resolve interindividual compositional differences based on cell surface protein information while alternative multiplexing methods can detect but not resolve encapsulation of multiple cells within a droplet.

Because pool-specific effects appear to be minimal in SCITO-seq (**Supplementary Fig. 13**), the pool-specific antibody barcodes could be used to directly label samples, obviating the need for orthogonal sample barcoding. To demonstrate this application, we stained cells from each of two donors in separate pools containing pool-specific barcoded antibodies (e.g., pool 1 contains CD4-BC1 staining donor 1 cells while pool 2 contains CD4-BC2 staining donor 2 cells). For loading concentrations of 2×10^4^ and 5×10^4^ cells, we obtained 17,730 and 34,549 post-processing CCD, sequenced to a per CCD depth of 964 and 1,540 reads for the ADT and 20,951 and 14,332 reads for the RNA. We observed the expected proportion of T and B cells per donor based on the distribution of the expression of CD4 and CD20 respectively (**Supplementary Fig. 14 and 15)**. After resolution, we recovered 18,680 and 41,059 cells at collision rates of 7.4% and 18.6% respectively (**Supplementary Fig. 16)**. Estimates of co-occurrence frequencies of different pool and antibody barcodes were highly correlated (r=0.99, *p*-value < 0.001) with observed values (**Supplementary Fig. 17**).

To demonstrate SCITO-seq’s applicability for high-dimensional and high-throughput cellular phenotyping, we profiled peripheral blood mononuclear cells (PBMCs) from two healthy donors using a panel of 28 monoclonal antibodies across 10 pools. After split pool staining, high concentration loading at 10^5^ and 2×10^5^ cells per microfluidic channel, and sequencing the resulting ADT and RNA libraries, we obtained 34,712 and 48,324 post-filtering CCDs. After resolution of the multiplets, we obtained collision rates of 4.4% and 8.5% and 55,420 and 93,127 resolved cells, increasing throughput by 10-fold over standard workflows. Leiden clustering based on either merged ADT counts or RNA across pools (**Fig. 3a**, **Supplementary Fig. 18** and **19**) resulted in clusters that are poorly differentiated in UMAP space due to the high multiplet rates at these loading concentrations. Strikingly, Leiden clustering using resolved ADT counts resulted in 17 clusters that are spatially differentiated in UMAP space and can each be annotated based on the expression of lineage specific markers (**Fig. 3b**). We detected seven clusters of monocytes, one cluster of conventional dendritic cells, naïve and memory CD4^+^ and CD8^+^ T cells, natural killer (NK) cells, B cells and gamma delta T cells (gdT). Notably, naive (CD45RA) and memory (CD45RO) CD4^+^ and CD8^+^ T cells emerge as distinct clusters which are often difficult to resolve using only transcriptomic data due to the low transcript expression of lineage (e.g. CD4) and state (e.g. CD45RO) determining markers^16^.

**Figure 3:**
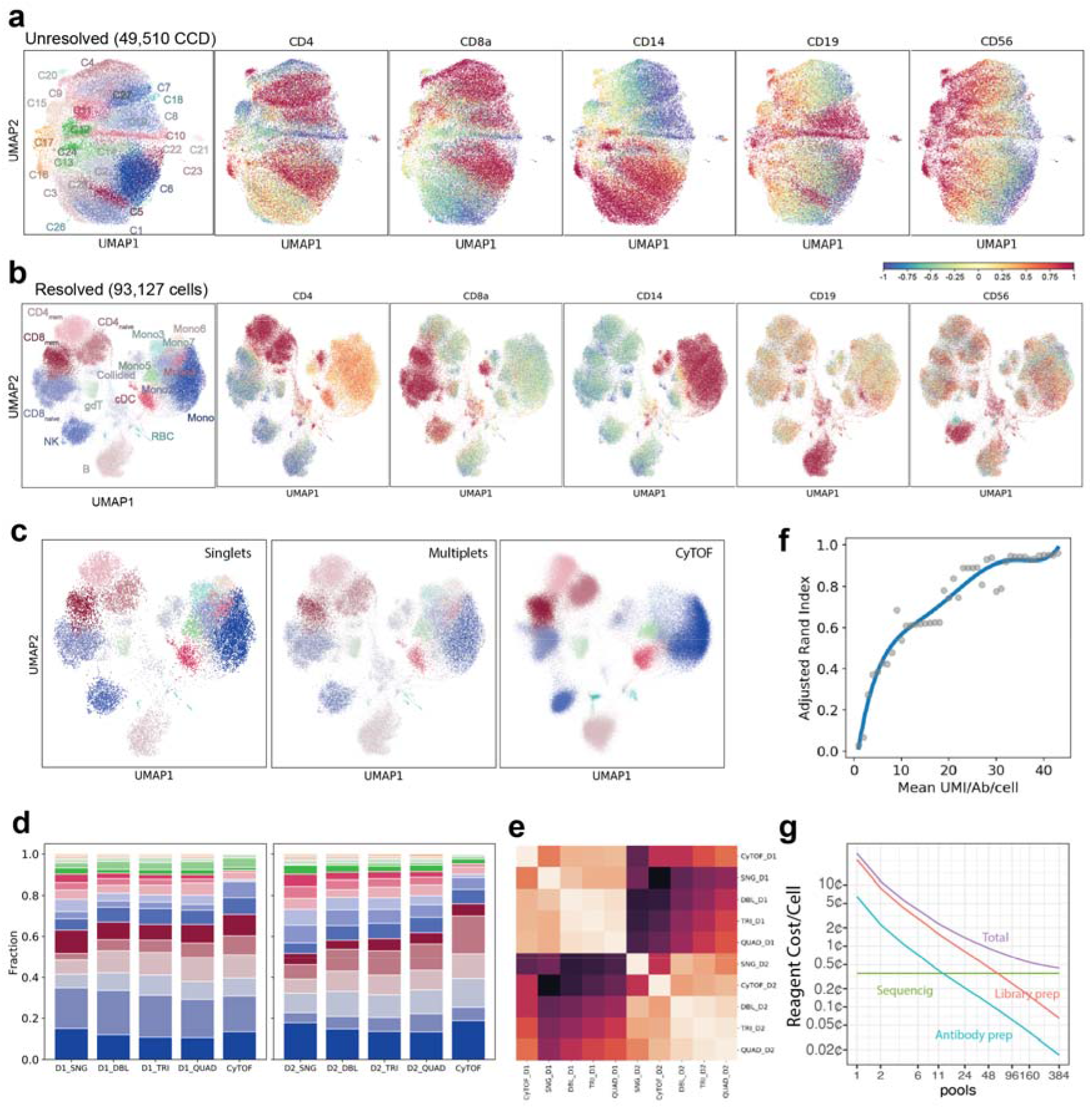
Large-scale PBMC profiling of healthy controls using antibody counts. **(a)** UMAP projection of unresolved droplet expression based on antibody counts showing key lineage markers when loading 2 ×10^5^ cells. **(b)** Ab+PBCs allow resolution of representative cell types in PBMCs utilizing an immunophenotyping panel of 28 cell surface antibodies. Representative markers displayed enrichment of ADT counts in specific cell clusters corresponding to canonical cell types in PBMCs. **(c)** UMAP projections of cells classified from pool singlets, multiplets and CyTOF. Principle Component Analysis (PCA)-based integration of data (Ingest function from Scanpy) was used to determine overlapping cell populations in SCITO-seq and CyTOF. **(d)** Stacked bar charts of cell type proportions among pool singlets and various multiplets (DBL: doublets, TRI: triplets, QUAD: quadruplets) and CyTOF within donor and across donors (D1 and D2). Each color maps to its respective Leiden cluster. **(e)** Heatmap of pairwise cosine similarity between estimated cell type proportions for singlets (SNG), doublets (DBL), triplets (TRI), quadruplets (QUAD) and CyTOF per donor. **(f)** Adjusted Rand Index of the resulting clusters (y-axis) versus down sampling of number of UMIs (per antibody per cell; x-axis). **(g)** Total cost estimates (purple) including library prep (red), antibody prep (blue) and sequencing cost assuming 45 reads/Ab/cell and a panel of 50 antibodies (green).

We further assessed the accuracy of SCITO-seq for quantitative immune phenotyping by comparing the compositional estimates obtained from pool singlets versus pool multiplets focusing only on droplets that contain cells from the same donor (as estimated using demuxlet). UMAP projections for cells resolved from pool singlets vs pool multiplets were qualitatively similar (**Fig. 3c**). The compositional estimates of the 16 immune populations detected from pool singlets and pool multiplets (doublet, triplet, quadruplets) from the same donor were highly similar as measured by cosine similarity (CS)(average CS over all pairwise comparisons: 0.98 [donor1], 0.97 [donor2]; **Fig. 3d** and **e**), demonstrating that SCITO-seq is capable of resolving cell surface expression profiles of cells within multiplets. Furthermore, the cosine similarity within donors was higher than between donors (average CS: 0.83) as expected. To further evaluate the data produced by SCITO-seq, we performed mass cytometry (CyTOF) using the same antibodies conjugated to metal isotopes (**Supplementary Fig. 20)**. Joint clustering of the CyTOF and SCITO-seq data produced qualitatively similar UMAP projections with comparable clusters (**Fig. 3c**) and quantitative comparisons of composition estimates per donor revealed similar distributions obtained from the two orthogonal assays (SCITO-seq vs CyTOF donor1 average CS: 0.95, SCITO-seq vs CyTOF donor2 average CS: 0.93) (**Fig. 3e** and **Supplementary Fig. 21).**

A key advantage of SCITO-seq as a tool for high-throughput phenotyping is the high information content obtained by profiling protein abundance at limited depth of sequencing per cell. This was demonstrated by down sampling to 25 UMIs/Ab/cell, which corresponds to ~45 reads/Ab/cell (at 45% library saturation), in the dataset generated from 2×10^5^ loading. At this sampling rate, we were able to achieve an Adjusted Rand Index (ARI) of > 0.8 for assigning cells to the same clusters in the full dataset (**Fig. 3f**). A similar trend was observed for the data generated from loading 10^5^ cells (**Supplementary Fig. 22**). For SCITO-seq, as the library preparation cost quickly diminish with increasing number of pools, the total cost per cell is dominated by sequencing. By shallow sequencing a limited number of targets (e.g. ~2×10^3^ reads per cell for ~50 targets), SCITO-seq can remain cost effective even when large numbers of pools are used (**Fig. 3g**). The cost-effective, simple and flexible design provide potential for incorporating additional modalities and orthogonal experimental information. This positions SCITO-seq as a compelling method for scalable rich phenotyping, especially for high-throughput screening and clinical biomarker profiling applications where targeted profiling of large numbers of cells across many samples are required.

## Methods

### Closed form derivation of collision and empty droplet rates

Suppose there are P pools of cells. For pool *p*, cells arrive according to a Poisson point process with rate *λ*_*p*_ >0 (abbreviated PPP(*λ*_*p*_)), where the unit of time corresponds to the inter-arrival time of droplets. In the most general formulation, we assume that the point processes for different pools are independent. Further, we assume the probabilities of a gel/bead and a cell encapsulated into a droplet as 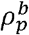 and 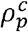, respectively. Therefore, by Poisson thinning, the arrival of cells follows 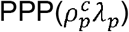.

We are interested in the probability of the event (called collision) that a droplet contains two or more cells from the same pool. Let *N*_*p*_ denote the number of cells from pool *p*, successfully loaded into a droplet. Then, *N*_1_, *N*_2_, ···, *N*_*P*_ where 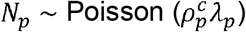, are independent random variables, and ℙ[Collision] can be computed as 1 − ℙ[No Collision]. Here ℙ[No Collision] represents a probability that every droplet contains ≤ 1 pool barcode. Therefore, we derive:

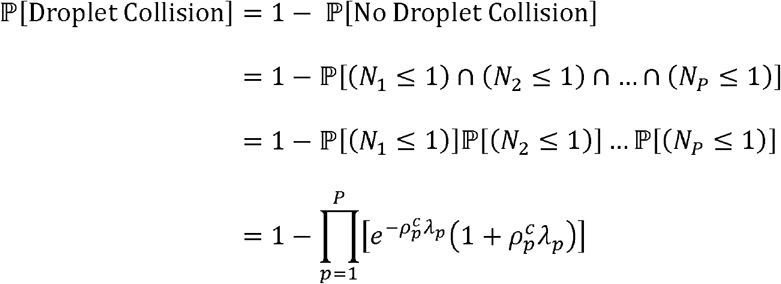

where the third equality follows from independence.

Next we condition ℙ[Droplet Collision] on ℙ[Non-empty Droplet], which is the probability that a droplet contains a cell at a given observation, ℙ[Non-empty Droplet] = 1 − ℙ[Empty Droplet], where:

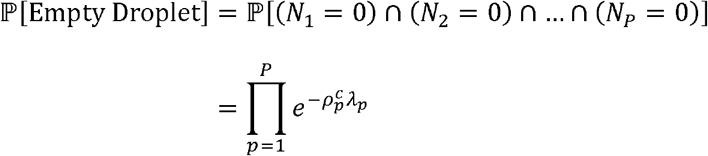

If there are *D* droplets formed and a total of *C* cells loaded evenly across the *P* pools (i.e., there are 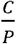 cells per pool), then 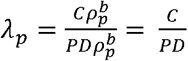 for all pools *p* = 1,2, …, *P* and that 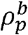 becomes a nuisance parameter. If we further assume that 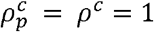 for all *p* = 1,2, … , *P*, then ℙ[Droplet Collision] and ℙ[Empty Droplet] simplify as

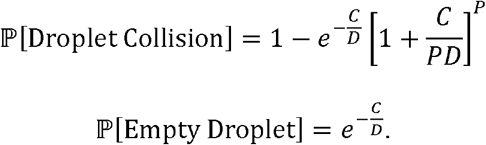

And finally, to estimated conditioned probability of barcode collisions:

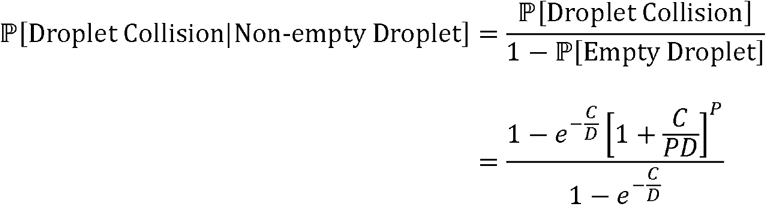

A second collision rate we can calculate is the cell barcoding (droplet barcode + pool barcode) collision rate which can be computed as the conditional probability that a particular pool *p* ∈ {1,2,… , *P*} has a collision in a given droplet, given that the droplet contains at least one cell from that pool. If we assume that there are *D* droplets formed and a total of *C* cells are distributed evenly across *P* pools, then we obtain:

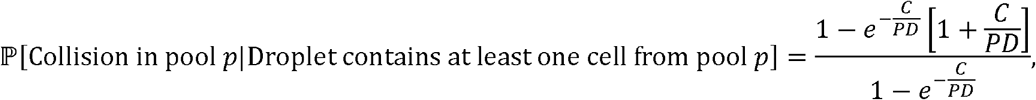

for all *p* ∈ {1,2, …, *P*}.

The above conditional probability is related to the proportion of the number of pools with a collision in a given droplet, relative to the total number of pools each with at least one cell represented in the droplet. More precisely,

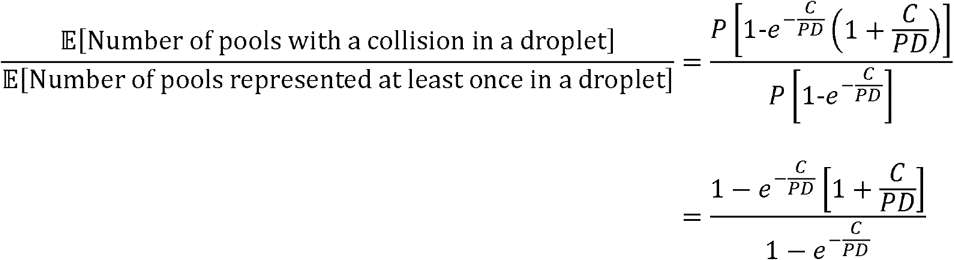

### Simulation of collision and empty droplet rate

For simulating the collision rates and empty droplet rates, we assumed a cell recovery rate of 60% and 10^5^ droplets are formed per microfluidic reaction resulting in *D* = 6 ∗ 10^4^. For *C* cells loaded, cell containing droplets are simulated using a Poisson process where λ = *C*/*D*. Assuming each simulated droplet *i* contains *γ*_*i*_ cells, we then compute the number of pool barcodes not tagging a cell in each droplet as:

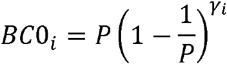

the number of pool barcodes tagging exactly one cell as:

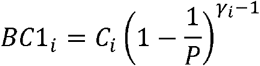

and the number of pool barcodes tagging greater than one cell as:

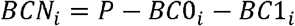

The conditional collision rate is estimated as:

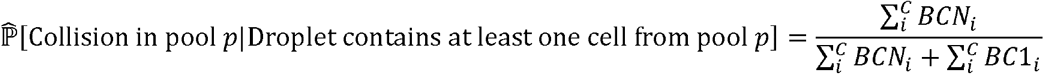

### Estimates of antibody conjugation, library construction, and sequencing

Cost for library conjugation is estimated to be $7.5 per antibody per *μ*g using the Thunderlink conjugation kit. Cost for library preparation is estimated to be $1,482 per well as advertised by 10X Genomics. Cost for sequencing is estimated as $26,700 per 12B reads as advertised by Illumina.

### Primary antibody oligonucleotide conjugation

For the species mixing experiment, anti-human CD29 and anti-mouse CD29 antibodies were purchased from Biolegend (cat. 303021, 102235) and conjugated per antibody using a ThunderLink kit (Expedeon cat. 425-0000) to distinct 20 bp 3’ amine-modified HPLC-purified oligonucleotides (IDT) to serve as hybridization handles. Antibodies were conjugated at a ratio of 1 antibody to 3 oligonucleotides (oligos). In parallel, oligos similar to current antibody sequencing tags were directly conjugated at the same ratio for comparison. Sequences for the hybridization oligonucleotides and directly conjugated oligos were designed to be compatible with the 10x feature barcoding system by introducing a reverse complementary sequence to the bead capture sequence, alongside a pool and antibody specific barcode for demultiplexing. Conjugates were quantified using Protein Qubit (Fisher cat. Q33211) for antibody titration and flow validation. Also, we orthogonally quantified the antibodies using protein BCA. For the human donor mixing experiment, CD4 and CD20 antibodies (Biolegend cat. 300541, 302343) were conjugated as described above.

### Antibody-specific hybridization design

After conjugation of primary handle oligos, antibodies were combined and pools of oligos (IDT) were used to hybridize the primary handle sequences prior to staining. Of note, only one conjugation was done per antibody with the previously mentioned 20 bp oligonucleotide (e.g. all CD4 conjugates have the same 20 bp oligonucleotide). To avoid non-specific transfer of oligonucleotides between the different antibody clones and the same antibody clone from different wells, each clone received a unique 20 bp handle (Antibody handle). To sequence with antibody and pool specificity, a 10 bp barcode was added to the secondary oligo. The total oligonucleotide sequence consisted of a reverse complementary sequence to the antibody specific primary handle sequence (20 bp), TruSeq Read2 (34 bp), pool barcode (10 bp), and capture sequence (22 bp) (**Fig. 1b**). Prior to cell staining, 1 ug of each antibody was pooled and hybridized with 1 ul of respective secondary oligonucleotides at 1 uM at room temperature for 15 minutes. The hybridized antibody-oligonucleotide conjugates were purified using an Amicon 50K MWCO column (Millipore cat. UFC505096) according to the manufacturer's instructions to remove excess free oligonucleotides.

### Determination of non-specific transfer of oligonucleotides between antibodies

To determine the optimal concentration of hybridizing oligonucleotides for cell staining, we performed a mixed cell line experiment to determine the level of background staining of free oligonucleotides. A mixture of lymphoblastoid cells and primary monocytes were stained with CD14 and CD20 antibodies and hybridized with oligonucleotides with different fluorophores (FAM and Cy5 respectively) per antibody for 15 minutes at room temperature. Concentrations of hybridizing oligonucleotides with different concentrations (1uM and 100 uM) were tested (**Supplementary Figure. 23**). Antibodies directly conjugated to fluorophores served as a positive control antibodies (CD13-BV421, Biolegend cat. 562596) to gate respective populations.

### Validation of saturation of hybridization oligonucleotides using flow cytometry

To determine the saturation of available primary oligo handles, 1 ug of conjugated CD3 antibody (Biolegend) was hybridized with 1 ul of 1 uM of a reverse complementary oligo with a Cy5 modification (IDT modification /5Cy5/). After incubating at room temperature for 15 minutes, 1 ul of 1 uM of the same reverse complementary oligo but with a FAM modification (IDT modification /56-FAM/) was added to the reaction and additionally incubated for 15 minutes. The cocktail was then added to 1×10^6^ PBMCs pre-stained with Trustain FcX (Biolegend cat. 422302) (**Supplementary Figure. 24**).

### 10x Genomics run for SCITO-seq

Washed and filtered cells were loaded into 10x Genomics V3 Single-Cell 3’ Feature Barcoding technology for Cell Surface Proteins workflow and processed according to the manufacturer’s protocol. After index PCR and final elution, all samples were run on the Agilent TapeStation High Sensitivity DNA chip (D5000, Agilent Technologies) to confirm the desired product size. A Qubit 3.0 dsDNA HS assay (ThermoFisher Scientific) was used to quantify final library for sequencing. Libraries were sequenced on a NovaSeq 6000 (Read1 28 cycles, index 8 cycles and Read2 98 cycles).

### Mixed species experiment

HeLa and 4T1 cells were ordered from ATCC (ATCC cat. CCL-2, CRL-2539) and cultured in complete DMEM (Fisher cat. 10566016,10% FBS (Fisher cat. 10083147) and 1% penicillin-streptomycin (Fisher cat. 15140122)) in a 37°C incubator with 5% CO2 on 10 cm culture dishes (Corning). Prior to staining, cells were trypsinized at 37°C for 5 minutes using 1 ml Trypsin-EDTA (Fisher cat. 25200056) and were quenched with 10 ml complete DMEM. Cells were harvested and centrifuged at 300xg for 5 minutes. Cells were resuspended in staining buffer (0.01% Tween-20, 2% BSA in PBS) and counted for concentration and viability using a Countess II (Fisher cat. AMQAX1000). HeLa and 4T1 cells were then mixed at equally and 1×10^6^ cells were aliquoted into two 5 ml FACS tubes (Falcon cat. 352052) and volume normalized to 85 ul. Cells were stained with 5 ul of Trustain FcX for 10 minutes on ice. Cell mixtures were stained with a pool of human and mouse CD29 antibodies, either with the direct or universal design, in a total of 100 ul for 45 minutes on ice. Cells were then washed 3 times with 2 ml staining buffer and centrifuged at 300xg for 5 minutes to aspirate supernatant. Cells were then resuspended in 200 ul of staining buffer and counted for concentration and viability as before. Cells from each stained pooled were mixed and 2×10^4^ or 1×10^5^ cells were loaded into the 10x chromium controller using 3’v3 chemistry.

### Human donor mixing experiment

PBMCs were collected from anonymized healthy donors and were isolated from apheresis residuals by Ficoll gradient. Cells were frozen in 10% DMSO in FBS and stored in a freezing container at −80°C for one day before long term storage in liquid nitrogen. Cells from two donors were quickly thawed in a 37°C water bath before being slowly diluted with complete RPMI1640 (Fisher cat.61870-036, supplemented with 10% FBS and 1% pen-strep) before centrifugation at 300xg for 5 minutes at room temperature. Cells were resuspended in EasySep Buffer (STEMCELL cat. 20144) at a concentration of 5×10^7^cells/ml before being subject to CD4 and CD20 negative isolation (STEMCELL cat. 17952, 17954). Isolated cells were counted and mixed at a ratio of 3 CD4:1 CD20 for donor 1 and a ratio of 1 CD4:3 CD20 for donor 2 for a total of 1.2×10^6^ cells per donor. The cells were centrifuged at 300xg for 5 minutes at room temperature and resuspended in 85 ul of staining buffer and incubated with 5 ul of Human TruStain FcX(Biolegend cat: 422301) for 10 minutes on ice in 5 ml FACS tubes. Cells from each donor were either mixed prior or stained with pool specific barcode hybridized antibody oligo conjugates for 30 minutes on ice. Staining was quenched with the addition of 2 ml staining buffer and washed as previously mentioned. Cells were resuspended in 0.04% BSA in PBS and cells from each well were counted, pooled equally, and then passed through a 40 um strainer (Scienceware cat. H13680-0040). The final strained pool was counted once more prior to loading into a 10x chip B with 2×10^4^ cells, 5×10^4^ cells, 1×10^5^ cells, and 2×10^5^ cells.

### Mass cytometry of healthy controls

PBMCs were isolated, cryopreserved, and thawed from the same donors as previously described. Once thawed, the cells were counted and 2×10^6^ cells from each donor were aliquoted into cluster tubes (Corning cat. CLS4401-960EA). Cells were live/dead stained with cisplatin (Sigma cat. P4394) at a final concentration of 5 uM for 5 minutes at room temperature. The live/dead stain was quenched and washed with autoMACS Running Buffer (Miltenyi Biotec cat. 130-091-221). Cells were then stained with 5 uL of TruStain FcX for 10 minutes on ice before surface staining. Mass cytometry antibodies were previously titrated using biological controls to achieve optimal signal to noise ratios. The antibodies in the panel were combined into a master cocktail and incubated with cells from the two donors and stained for 30 minutes at 4°C. After washing twice with 1 ml autoMACS Running Buffer, the cells were resuspended and fixed in 1.6% PFA (EMS cat. 15710) in MaxPar PBS (Fluidigm cat. 201058) for 10 minutes at room temperature with gentle agitation on an orbital shaker. Samples were then washed twice in autoMACs Running Buffer, and then three times with 1X MaxPar Barcode Perm Buffer (Fluidigm cat. 201057). Each sample was then stained with a unique combination of three purified Palladium isotopes obtained from Matthew Spitzer and the UCSF Flow Cytometry Core for 20 minutes at room temperature with agitation as previously described^26^. After three washes with autoMACS Running Buffer, samples were combined into one tube and stained with a dilution of 500 uM Cell-ID Intercalator (Fluidigm cat. 201057), to a final concentration of 300 nM in 1.6% PFA in MaxPar PBS at 4°C until data collection on the CyTOF three days later. Immediately before running on the CyTOF machine, the sample tube was washed once with each autoMACS Running Buffer, MaxPar PBS, and MilliQ H2O. Once all excess proteins and salts were washed out, the sample was diluted in Four Element EQ Calibration Beads (Fluidigm cat. 201078) and MilliQ H2O to a concentration of 1e6 cells/mL and run on a CyTOF Helios at the UCSF Flow Cytometry Core.

### Comparing Mass cytometry (CyTOF) and SCITO-seq

Data was transferred from the CyTOF computer and normalized and de-barcoded using the premessa package (https://github.com/ParkerICI/premessa). Clean files were uploaded to Cytobank (https://www.ucsf.cytobank.org/) for gating and manual identification of immune cell subsets. Files containing only singlet events were exported from Cytobank and analyzed with CyTOFKit2 package (https://github.com/JinmiaoChenLab/cytofkit2). Through CyTOFkit2, events were clustered using Rphenograph with k=150 and visualized via UMAP for proportion determination.

### Pre-processing and initial filtering

Both the species mixing experiments and human donor mixing experiments were processed using Cell Ranger 3.0 Feature Barcoding Analysis using default parameters. For cDNA and ADT alignment, we specified the input library type as ‘Gene Expression’ and ‘Antibody Capture’ respectively as recommended. For ADT alignment, specific barcode sequences (Ab+pool) were specified as a reference. Reads were aligned to the hg19 and mm10 concatenation reference for species mixing experiment. For all human experiments, the reads were aligned to the human reference genome (GRCh38/hg20). We first removed RBC and Platelets and removed cells with more than 15% of mitochondrial gene related reads. We further removed genes with less than 1 counts across all cells.

### Normalization for species mixing and T/B cell human donor mixing experiment

For cDNA counts, data was normalized by dividing each UMI counts to the total UMI counts and multiplied by 10,000. Then, the data was log1p transformed (numpy.log1p). Finally, the data was scaled to have mean = 0 and standard deviation = 1. Clustering was done using the Leiden algorithm^27^ using 10 nearest neighbors and a resolution of 0.2 for mixed species and two-donor experiment with two cell types (T and B cells).

To normalize ADT counts in species mixing experiment, the data was log transformed and standardized to have mean = 0 and standard deviation = 1. For ADT counts in two human donor mixing experiment with two cell types, after log transformation of the raw data, we used a Gaussian Mixture Model in scikit-learn package in python to normalize the data with the following parameters (convergence threshold 1e-3 and max iteration to 100, number of components 2). The data was normalized by z-score like transformation (log transformed raw value - mean of the posterior means of two components / mean of the posterior standard deviations).

### Implementation of an algorithm for pool demultiplexing and multiplet resolution

Considering all antibodies in each pool, we normalized each value by dividing mean expression value of CD45 counts across all pool (considered as a universal expression marker) for each droplet barcode yielding a p*m matrix (p is the number of pool and m is number of droplet barcodes). Then, the matrix was CLR normalized and demultiplexed using HTODemux from Seurat (v3.0) (http://satijalab.org/seurat/) to classify the droplet barcode to a pool or unassigned (we discretized the value of 0 or 1). Using this binary matrix, we iterated over p times (where discretized value equals 1) to get final resolved matrix of (n*r) where n is the number of antibodies used and r is the resolved number of cells. For each iteration, we selected the columns that were positive for the above-mentioned discretized matrix. An additional round of HTODemux was used to re-classify the ‘Negative’ cells from initial classification because most of the initial classification which deemed the cells negative had a UMAP distributions which were contained in the original clusters.

### Analysis of PBMC experiment

#### Normalization and resolution of multiplets

To normalize cDNA data for PBMC experiments, we used the same normalization method as described above. To generate the UMAP based on ADT counts for the PBMC experiment, we performed pool demultiplexing using the algorithm described previously. Then, the resolved matrix (n*r) was normalized as in the cDNA processing. Raw values were normalized to total counts of 10,000 per cell and log1p transformed. Then, the values were standardized (mean 0, standard deviation 1) per pool. Using these normalized values, PCA was performed to reduce dimensionality. Leiden clustering was done with 10 neighbors and 15 PCs from the previous step. A resolution value of 1.0 was used to assign clusters for the whole PBMC experiments. Finally, UMAP was utilized to visualize the resolved total cells.

#### Demultiplexing donor identity

For demultiplexing the donors, a VCF file containing donor genotype information and the bam file output from the Cell Ranger pipeline were used as inputs for demuxlet with default parameters. For donors without genotypic information, we used vireo^23^ to assign droplet barcodes to the corresponding donor.

#### Downsampling experiment with Adjusted Rand Index calculations

To evaluate the quality of clustering at a given downsample depth, Adjusted Rand Index (ARI) was used as the representative metric. Leiden clustering was performed on the full dataset of RNA and ADT. Then, resulting cluster labels were taken as ground truth cell type assignments. To determine an optimal Leiden resolution for downsampling, clustering was performed 5 times at a range of resolutions. A resolution that produced consistently high ARI was then used to generate ground truth labels and perform clustering on downsampled data. Data was downsampled to a specified mean UMI/Antibody/cell using scanpy (1.4.5.post3) to downsample total reads. Downsampled data was then clustered and labels compared to full dataset clustering with ARI.

## Acknowledgments

C. J. Y. is a Chan Zuckerberg Biohub Intercampus Research Award Investigator. This work was supported by NIH grant DP5 OD023056 and funding from the UCSF Program for Breakthrough Biomedical Research to M.H.S. and NIH grant S10 1S10OD018040, which enabled the procurement of the CyTOF mass cytometer used in this study.

## Code availability

All code used to perform simulation and generating figures can be found at https://github.com/yelabucsf/SCITO-seq.

