## Supplementary Materials for "SCITO-seq: single-cell combinatorial indexed cytometry sequencing"

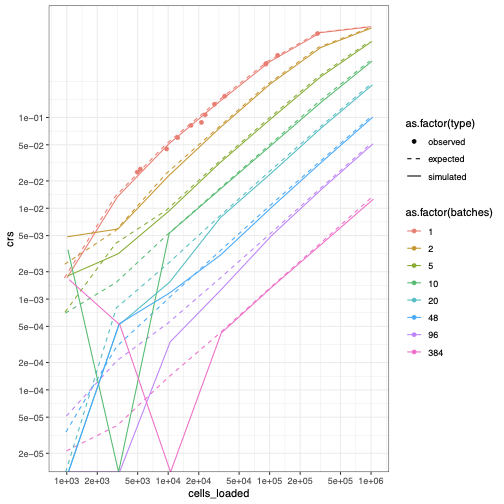

**Supplementary Figure 1.** Collision rate (y-axis, denoted as crs) as a function of cell loading concentration (x-axis, denoted cells_loaded) and number of pools (colored lines). Observed collision rates (points) are based on all experiments performed. Expected (dotted line) and simulated (solid line) collision rates are based on Poisson statistics where lambda = cells loaded / 100000 droplets and number of total cells containing droplets is modeled as 0.6 x 100000.

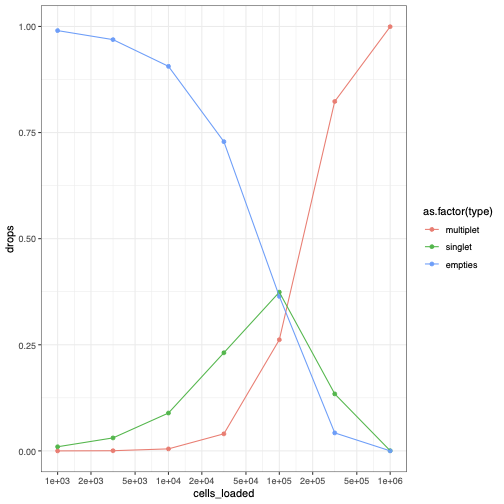

**Supplementary Figure 2.** Number of droplets (y-axis) containing no cells (blue), exactly one cell (green) or greater than one cell (red) as a function of the number of cells loaded (x-axis).

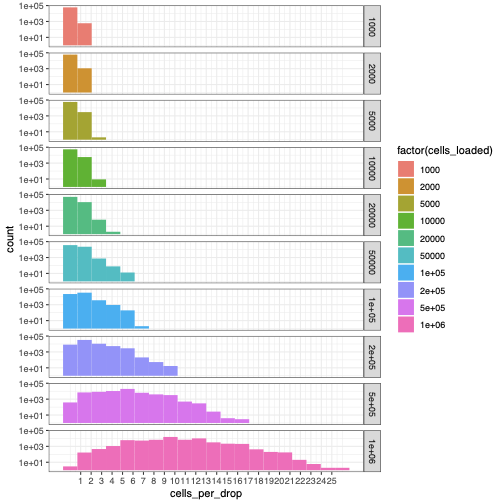

**Supplementary Figure 3.** Distribution of number of cells per droplet for different cell loading numbers (cells_loaded).

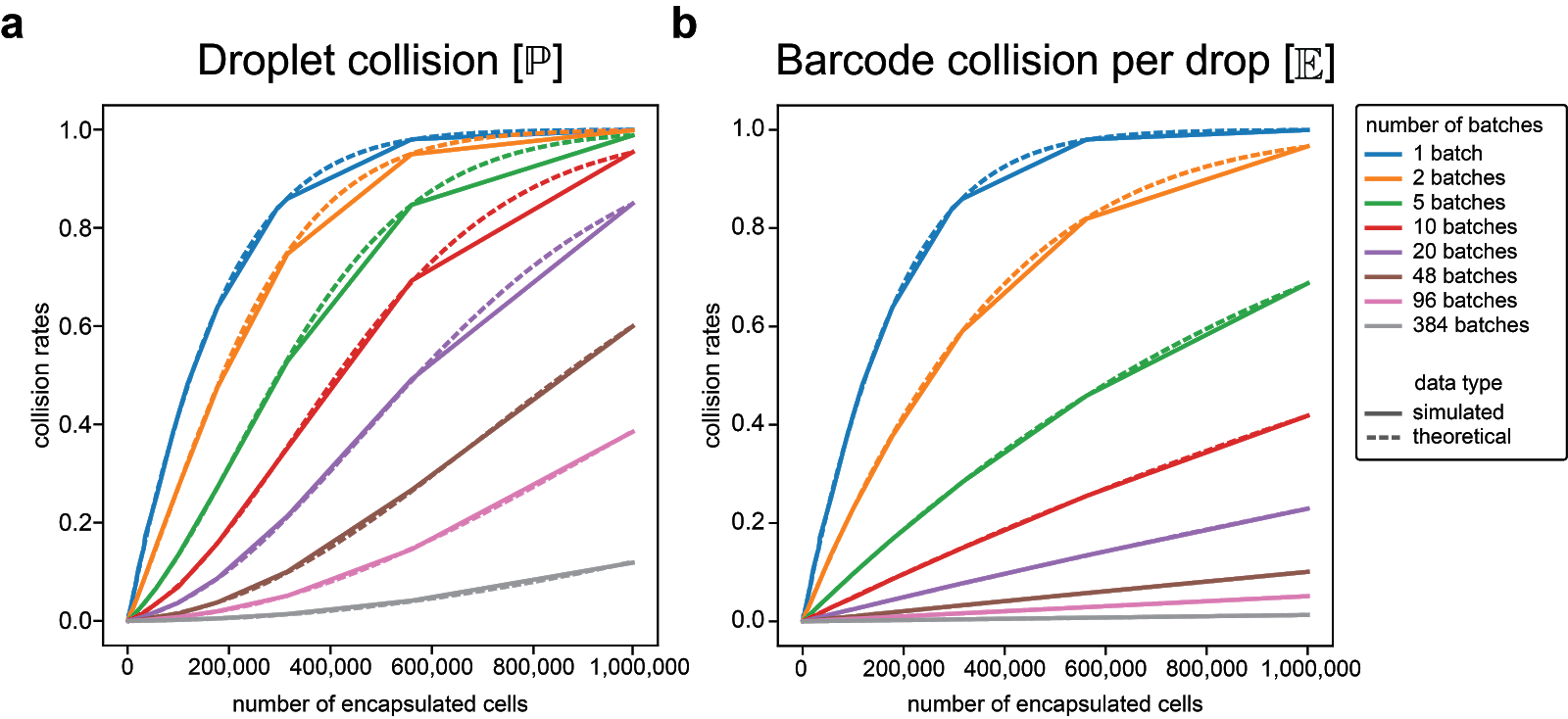

**Supplementary Figure 4. Comparison of the theoretical and simulated collision rates of SCITO-seq. a.** Droplet collision rate, depicting proportion of droplets with at least one barcode collision. **b.** Barcode collision rate, estimating proportion of batches (pools) with a collision in a given droplet. Collision rates were calculated using simulations of a Poisson Point Process (solid lines) or a closed form solution (dashed lines; see **Methods**). Estimates from a closed form solution robustly and almost identically recapitulate simulations and can be used to calculate collision rates for an experiment.

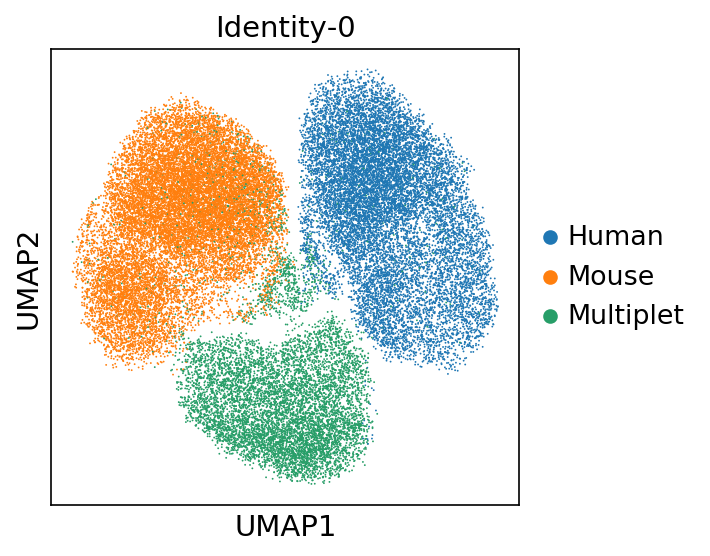

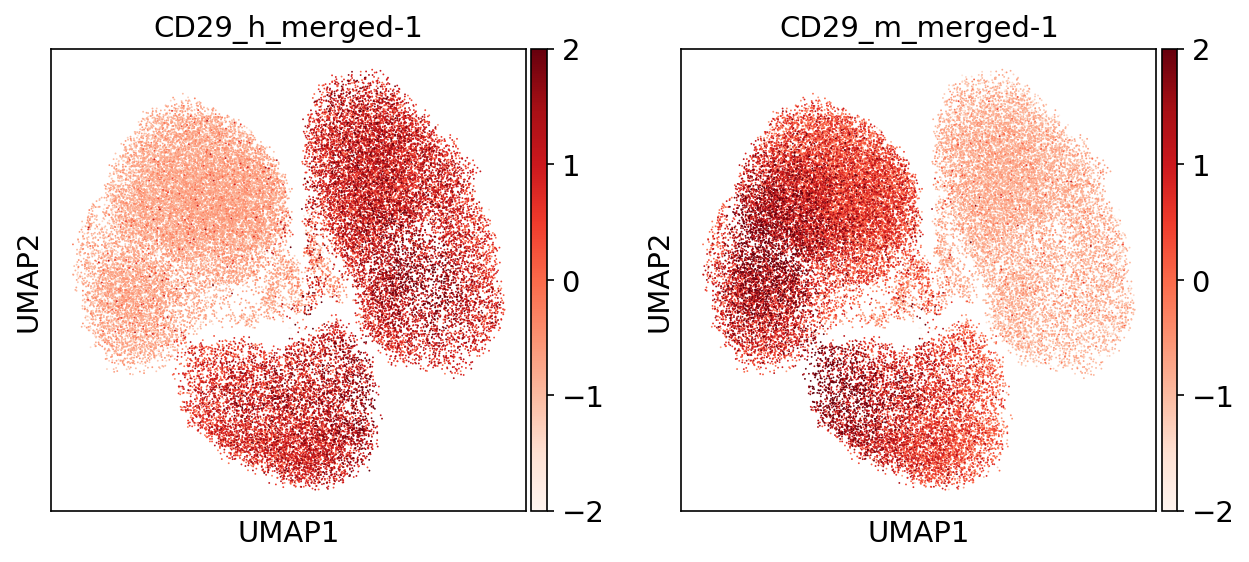

**Supplementary Figure 5.** Staining of mouse and human cells on transcriptome UMAP for 100k loading experiment. Upper figure panel color is based on transcriptome and for bottom plots, we merged two different pool barcodes per species (for example, CD29_h_merged-1 = CD29_h_barcode-1+....+CD29_h_barcode-5) .

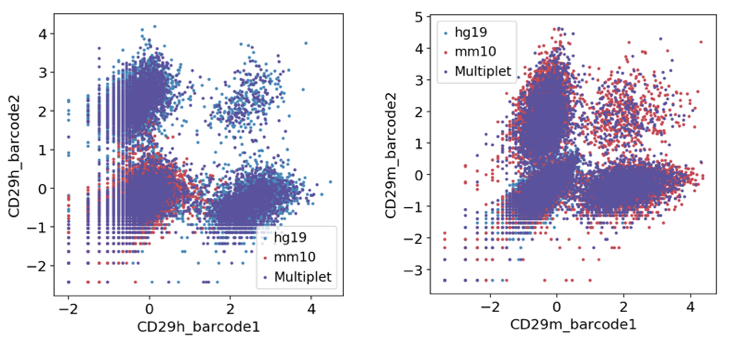

**Supplementary Figure 6.** Scatterplot showing within species doublet across batches for 100k loading species mixing experiment. We can resolve the double positive population (positive for both pool barcodes, such as CD29h_barcode1 and CD29h_barcode2 positive or CD29m_barcode1 and CD29m_barcode2 positive shown here).

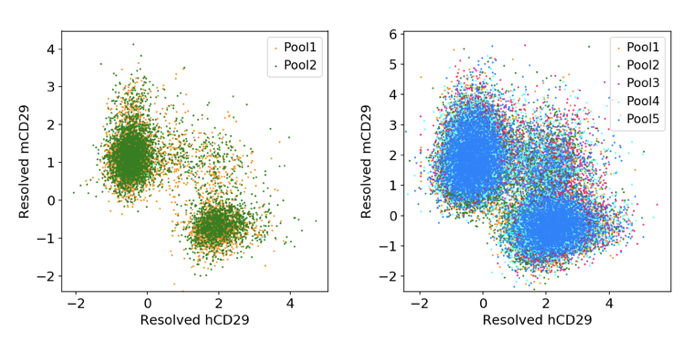

**Supplementary Figure 7.** Scatterplots for species mixing 20k (left) and 100k (right) loading experiments colored by pool showing batch specific staining level.

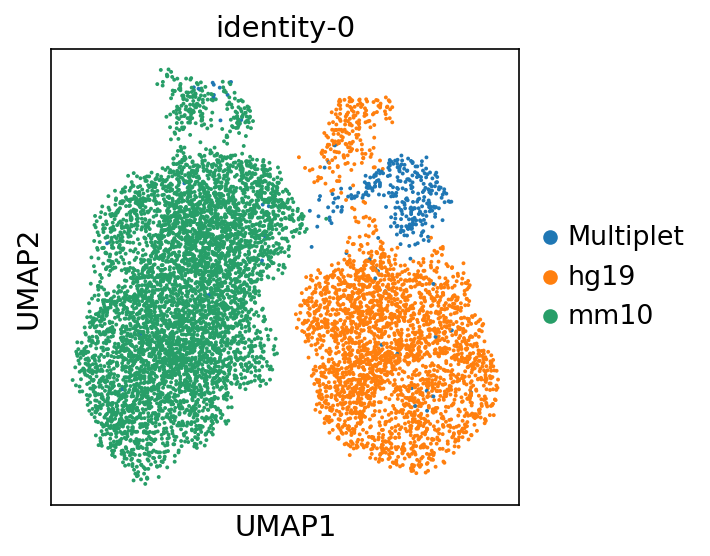

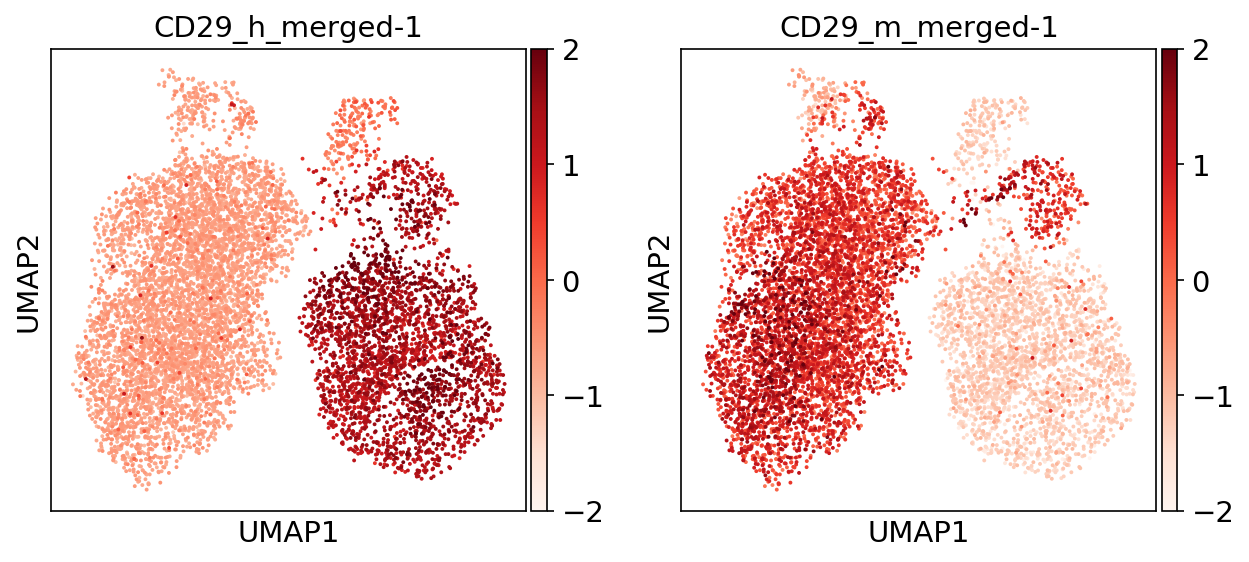

**Supplementary Figure 8.** Staining of mouse and human cells on transcriptome UMAP for 20k loading experiment. Upper figure panel color is based on transcriptome and for bottom plots, we merged two different pool barcodes per species (for example, CD29_h_merged-1 = CD29_h_barcode1+CD29_h_barcode2) .

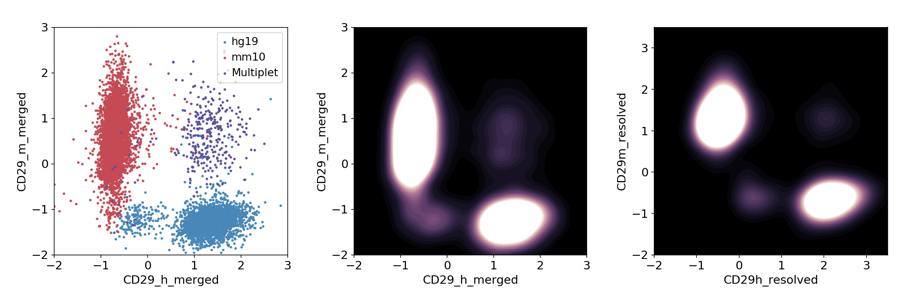

**Supplementary Figure 9.** Scatterplot (left, unresolved) and density plots (middle: unresolved, right: resolved) for 20k loading species mixing experiment.

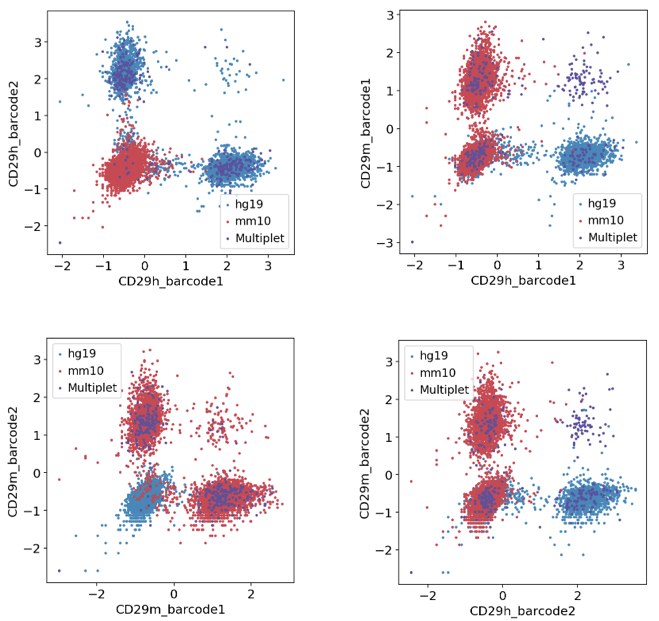

**Supplementary Figure 10.** Scatter plots of 20k loading of species mixing experiment to show cross pool or within pool background level using SCITO-seq.

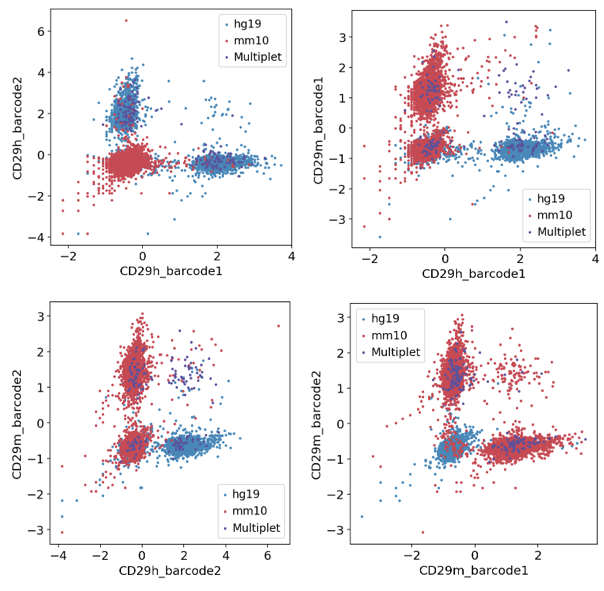

**Supplementary Figure 11.** Scatter plots of 20k loading of species mixing experiment to show cross pool or within pool background level using direct conjugation. All four possible comparisons are shown above.

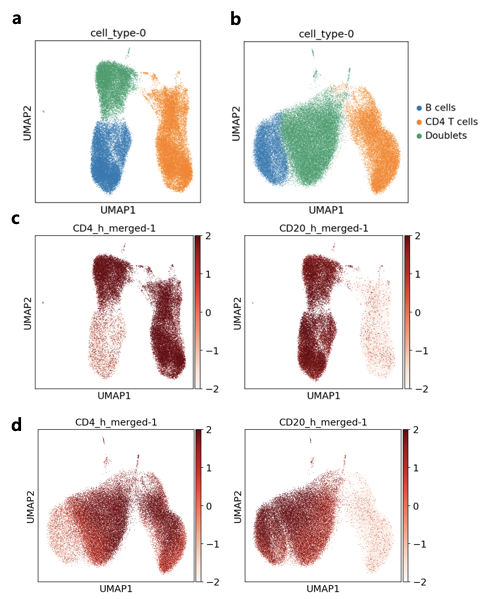

**Supplementary Figure 12.** UMAP projections of T/B cell experiments with (a) 100k and (b) 200k loading colored according to the cell types. The potential cell type ‘Doublets’ are also colored in green. The Bottom figure shows the staining level of merged SCITO-seq antibodies for (c) 100k and (d) 200k on top of RNA UMAP space.

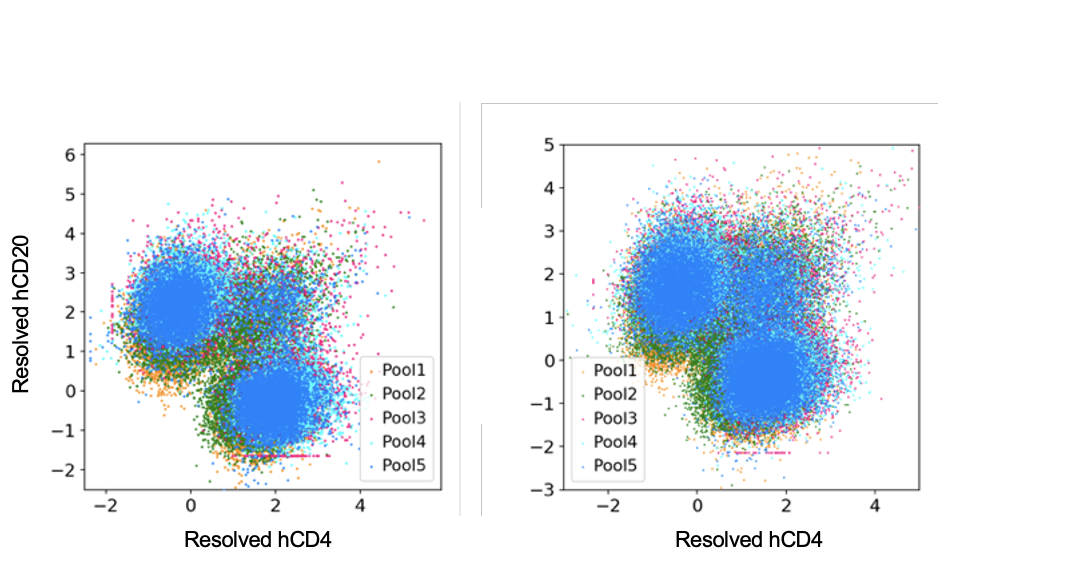

**Supplementary Figure 13.**  Scatterplots for species mixing 100k and 200k T/B experiments loading experiments colored by pool showing batch effects.

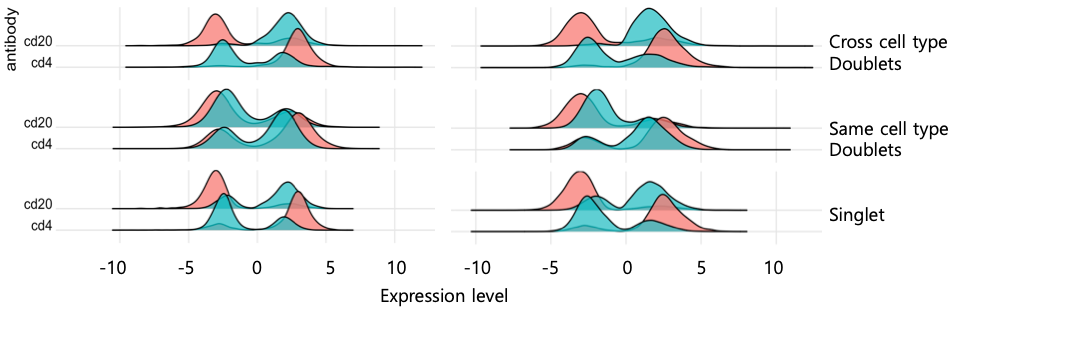

**Supplementary Figure 14.** Distribution of the normalized UMI counts for each antibody in cells resolved from pool singlets and pool multiplets per donor for 20k (left) and 50k (right) T/B mixture experiment (colors indicate different pools, pool1:red and pool2:green).

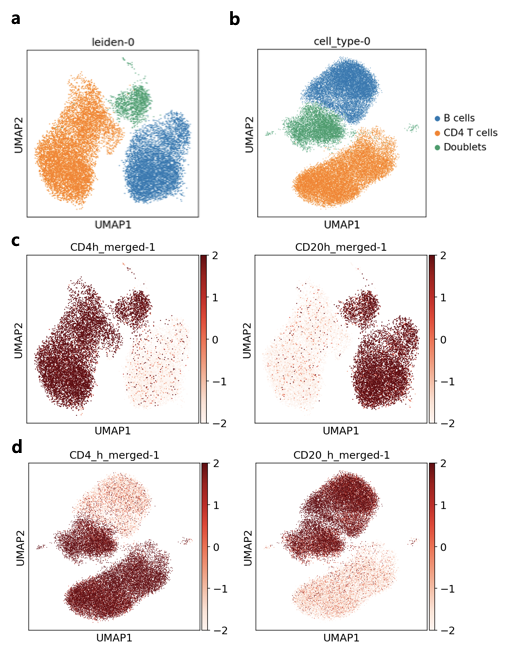

**Supplementary Figure 15.** UMAP projections of T/B cell experiments with (a) 20k and (b) 50k loading colored according to the cell types. The potential cell type ‘Doublets’ are also colored in green. The Bottom figure shows the staining level of merged SCITO-seq antibodies for (c) 20k and (d) 50k on top of RNA UMAP space.

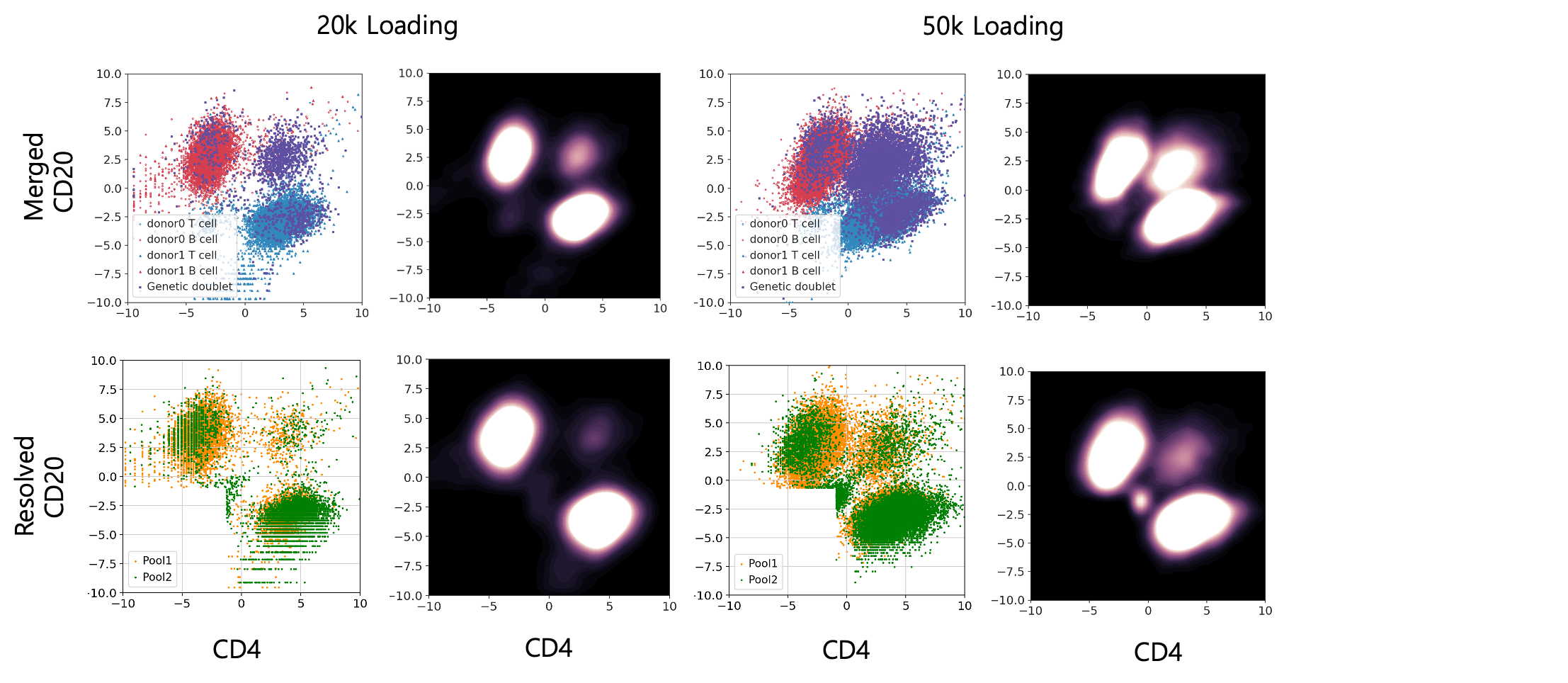

**Supplementary Figure 16.** Comparing the resolution of multiplets in 20k and 50k loading of T and B cell experiments. In this experiment, we stained two pools of antibodies with each pool stained with different donor cells.

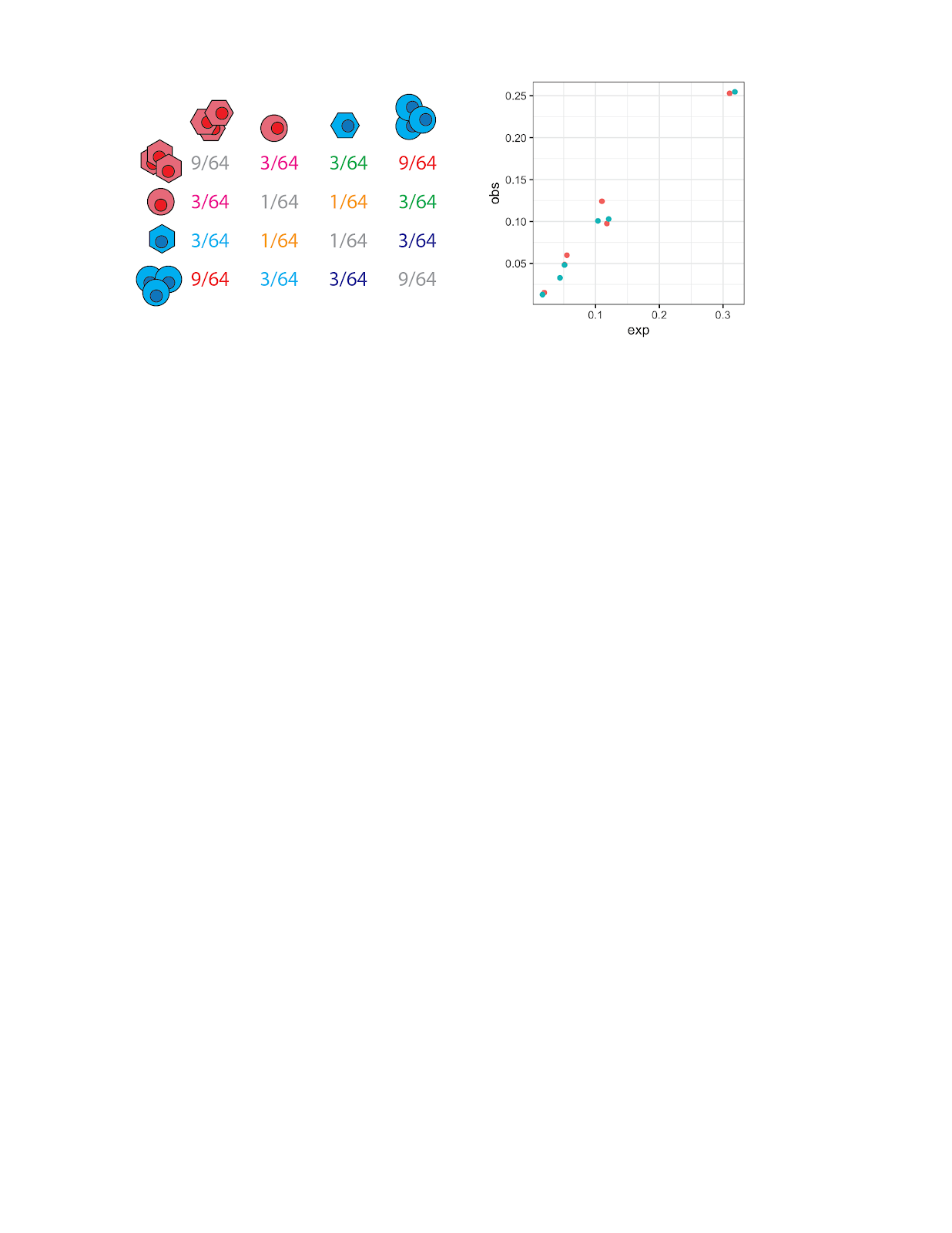

**Supplementary Figure 17.** Expected (x-axis) vs observed (y-axis) co-occurrence frequencies in 20k (green) and 50k (red) T and B cell loading experiments. Left matrix shows the expected particular cell-type multiplet probabilities (color scheme corresponds to the main Fig 2a.

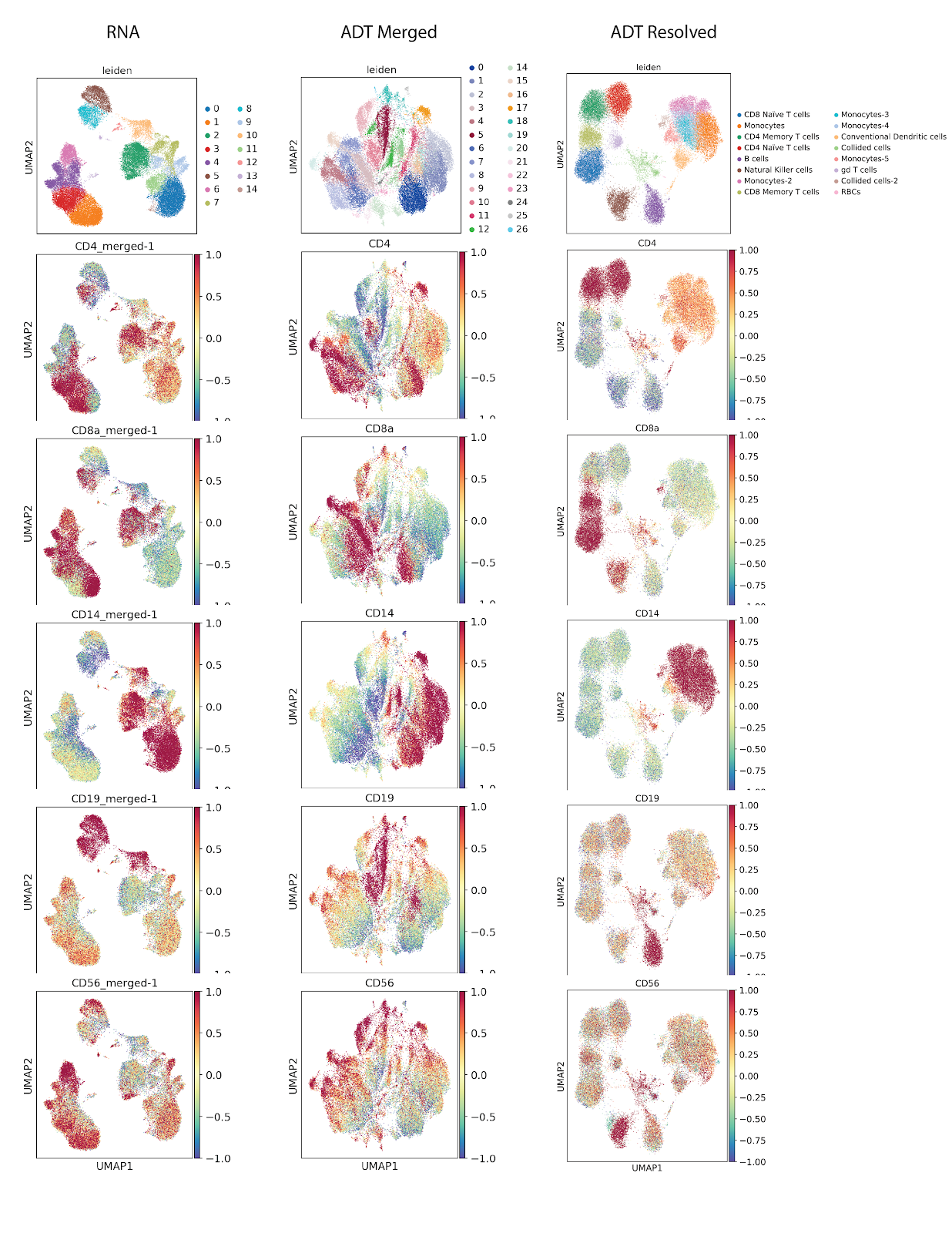

**Supplementary Figure 18.** UMAP projections for 100k PBMC loading. UMAP projection using RNA expression (left), merged antibody counts before resolution (middle) and after resolution (right).

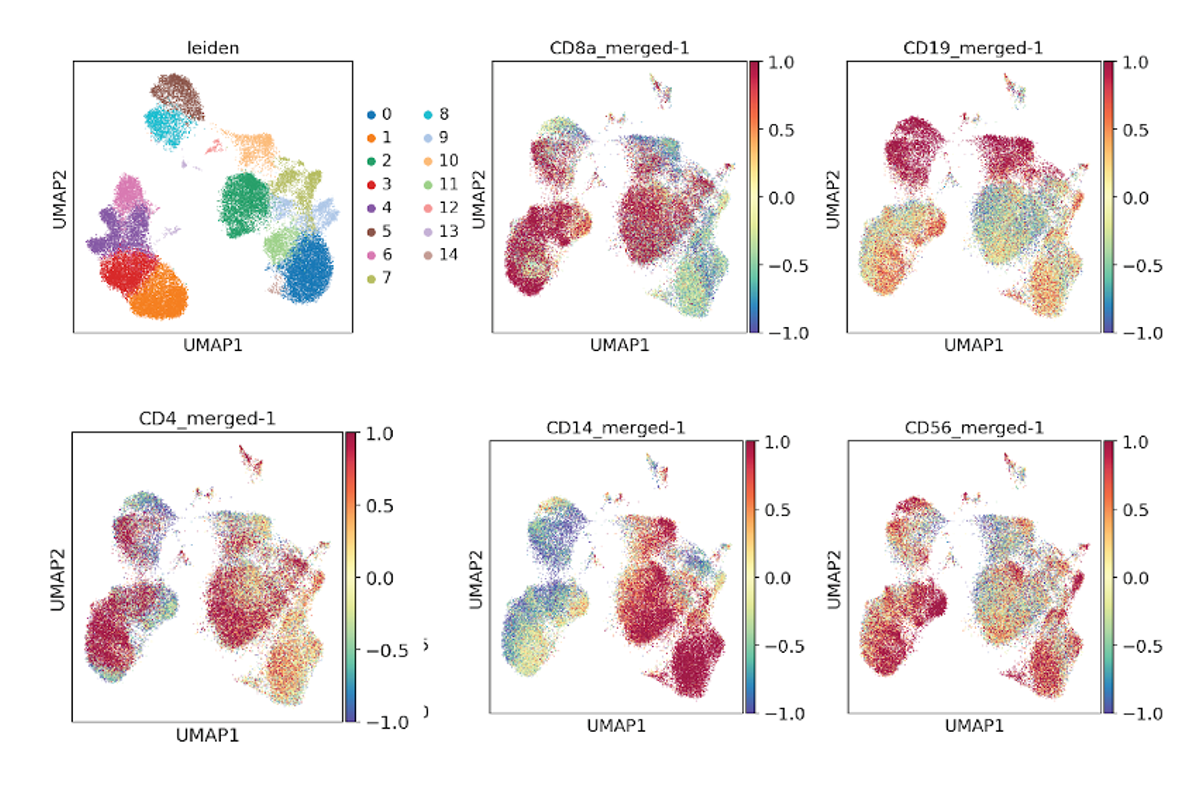

**Supplementary Fig 19.** UMAP RNA projections for representative markers of 200k PBMC loading.

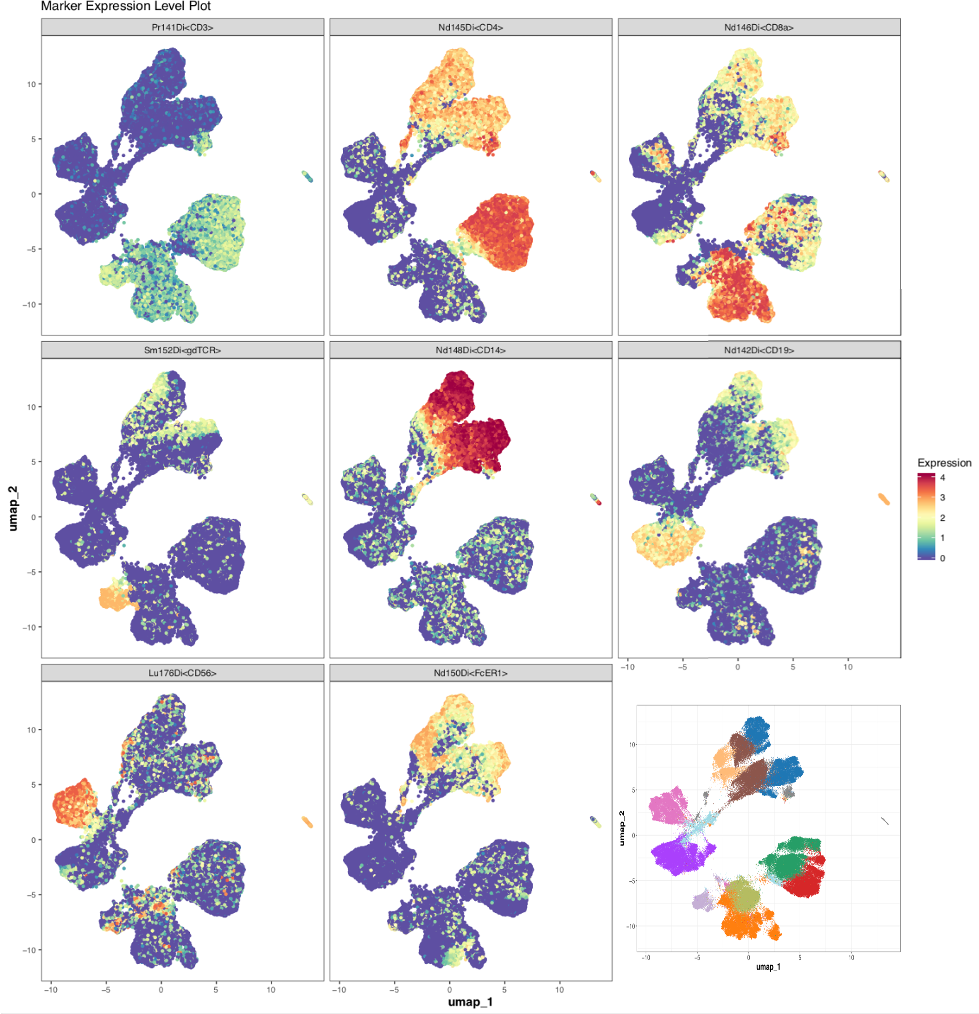

**Supplementary Figure 20.** UMAP with representative markers based on CyTOF experiment using the same antibody panel as in SCITO-seq.

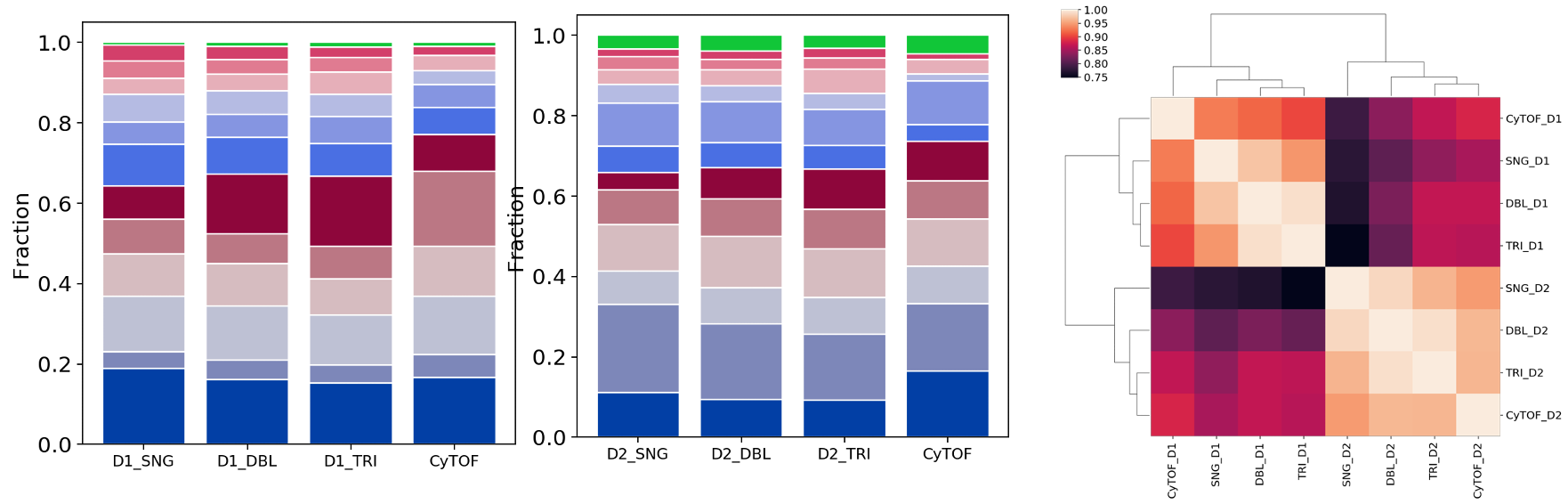

**Supplementary Figure 21.** Comparisons of SCITO-seq with CyTOF per donor (D1: left, D2: middle) for 100k data. Pairwise correlation heatmap plot (right) is also shown (similar to Fig3. e).

Within donors, the proportion each Leiden cluster was highly correlated (Cosine similarity within donor1:0.95, donor2:0.94).

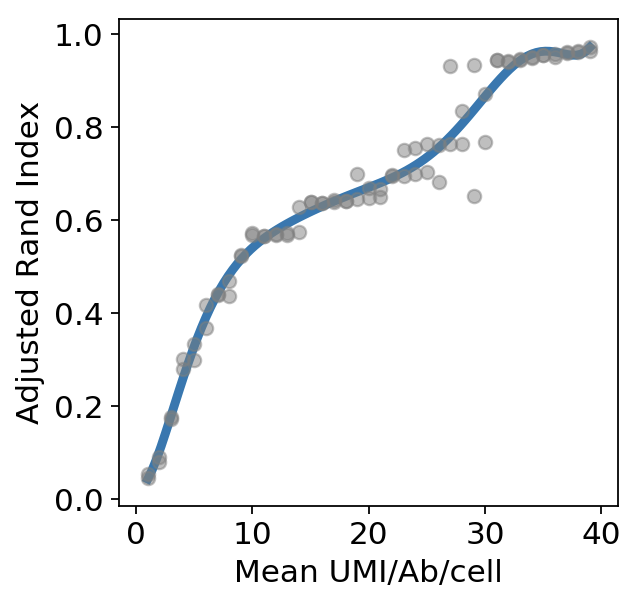

**Supplementary Figure 22.** Downsampling analysis of 100k loading data for Rand index analysis

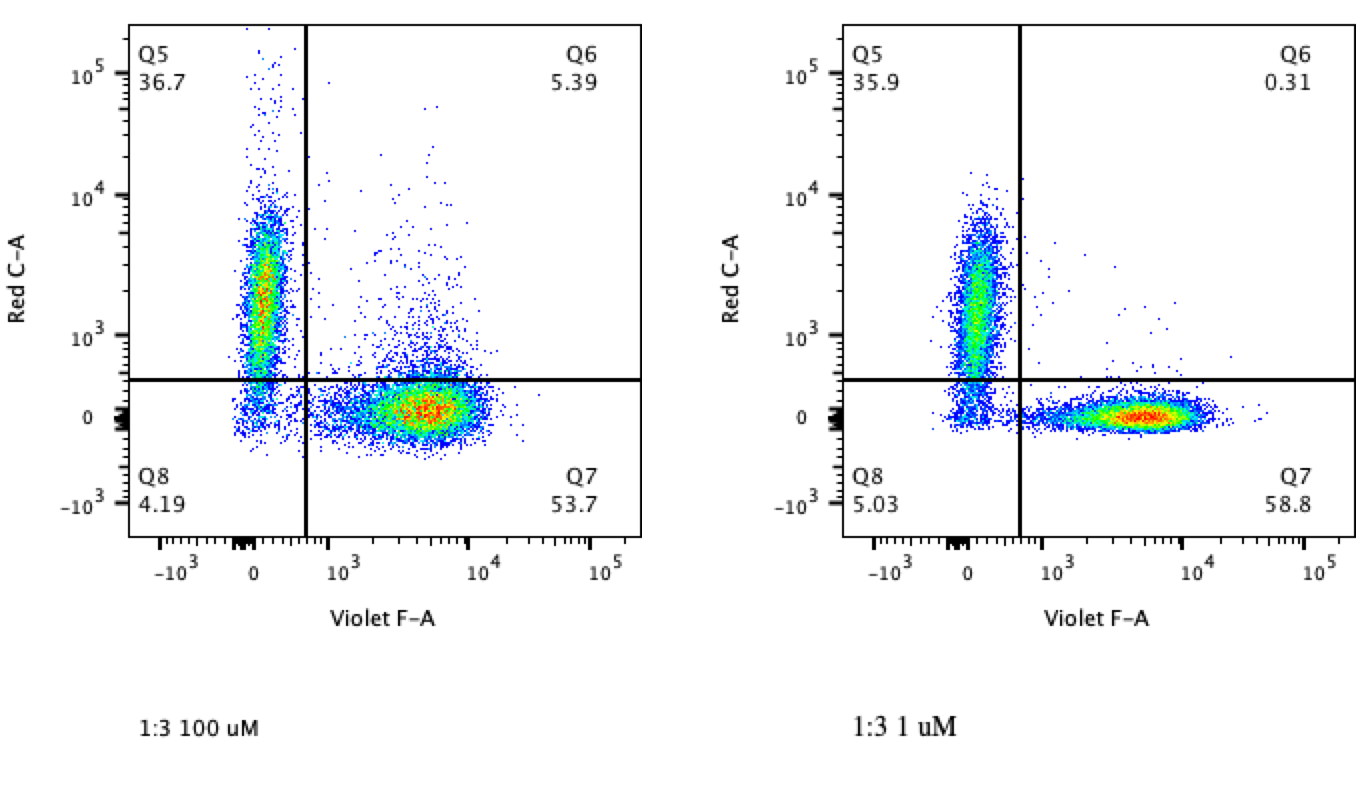

**Supplementary Figure 23. Titration experiment using Flow Cytometry**

To reduce the non-specific staining of secondary oligonucleotides, we titrated oligonucleotides at 1uM and 100uM. After hybridization of oligonucleotide conjugated antibodies with a Cy5 conjugated reverse complementary oligonucleotide for 15 minutes, a mixture of LCLs and primary monocytes were stained with the hybridized material an CD13-BV421 for 30 minutes, washed twice and analyzed on a LSRII. CD13 BV421 antibody was captured by the Violet-F channel (x-axis)  and Cy5 tagged secondary oligonucleotides was captured on the Red-C channel to check the level of background staining (Q6 gated population refers to the spillover of non-cognate secondary oligonucleotides in the primary monocyte population).

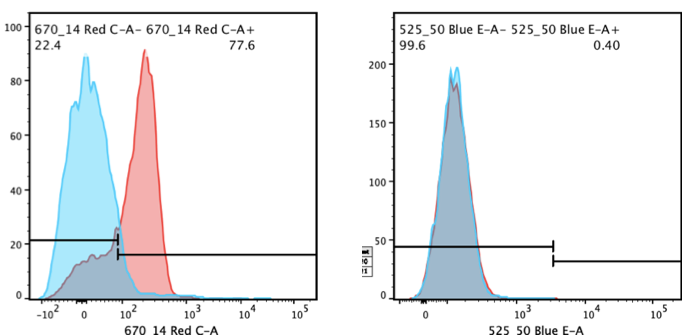

**Supplementary Figure 24. Saturation analysis using Flow Cytometry**

To determine if 1 ul of 1 uM reverse complementary oligonucleotide would saturate 1 ug antibody, we first hybridized 1 ug of oligonucleotide conjugated CD3 with 1 ul of 1 uM reverse complementary oligonucleotide conjugated to Cy5. Following this, another 1 ul of 1 uM reverse complementary oligonucleotide was added, but with a FAM conjugated instead. This was incubated for 15 minutes before being added to the whole PBMC and washed twice before running on an LSRII. Left figure shows the positive shift (red) when first hybridization occurs and second histograms shows there is essentially no significant shift because of the near saturation of the first handle sequence.
